# A cell death-independent role for MLKL in TGF-β receptor recycling in Marfan syndrome

**DOI:** 10.64898/2026.10.09.757966

**Authors:** Harshal Nemade, Henning Guthoff, Oscar Zimmermann, Sophia Riemen, Helena Stiller, Manuel Hedbawny, Bill Agbowo, Daniel Zahnow, Falk Holtkamp, Per Arkenberg, Dennis Mehrkens, Philipp Jahn, Felix Sebastian Nettersheim, Patrik Schelemei, Felix Ruben Picard, Kristel Martínez Lagunas, Jayashree Jayachandran, Gerhard Sengle, Uris Ros, Holger Winkels, Thomas Riffelmacher, Stephan Baldus, Ana J García-Sáez, Matti Adam, Martin Mollenhauer

## Abstract

Marfan syndrome (MFS) causes thoracic aortic aneurysm through mutations in the fibrillin-1 gene and subsequent dysregulated TGF-β signalling. Mixed lineage kinase domain-like protein (MLKL) executes necroptosis after RIPK3-dependent phosphorylation, but also functions in endosomal trafficking. Here we show that MLKL phosphorylation sustains TGF-β receptor recycling and aortic wall integrity without causing cell death. Phosphorylation-dead MLKL^S345A/S347A^ (*Mlkl*^AA^) mice developed aortic root dilation with disorganised collagen architecture and a secretory-pathway proteomic signature. Crossing them with *Fbn1^C1039G^*^/+^ MFS mice aggravated aortic dilation, disrupted collagen fibril ultrastructure and prolyl/lysyl hydroxylation, and caused lethal rupture under angiotensin II. Complete MLKL deficiency reproduced this phenotype. Phosphorylated MLKL was increased in MFS aortas, yet cleaved caspase-3, TUNEL positivity, smooth muscle cell content and leukocyte infiltration were unchanged. In smooth muscle cells, MLKL depletion dysregulated collagen-modifying enzymes, altered TGF-β-driven matrix remodelling and impaired RAB11-dependent TGFβR1 recycling. These findings suggest that phosphorylation-dependent, non-lytic functions of MLKL may contribute to the maintenance of thoracic aortic integrity in MFS.

## Introduction

Marfan syndrome (MFS) is the prototypical autosomal dominant connective tissue disorder, with a prevalence of approximately 1:3,000–1:5,000.^1^ It is caused by mutations in the extracellular matrix (ECM) protein fibrillin-1 (FBN1), leading to defective or absent microfibrils and impaired structural scaffolding of the aortic wall. This predisposes to thoracic aortic aneurysm (TAA), with a risk of dissection or rupture that remains the leading cause of premature death in affected individuals.^2^ Beyond their mechanical role, intact microfibrils sequester latent transforming growth factor-β (TGF-β) complexes within the matrix. In MFS, defective microfibrils fail to regulate TGF-β bioavailability, producing dysregulated canonical (SMAD-dependent) and non-canonical (MAPK/ERK) signalling that promotes ECM degradation, matrix metalloproteinase expression, smooth muscle cell (SMC) dysfunction and progressive weakening of the aortic wall.^3^

How TGF-β signalling is sustained in the MFS aorta is not explained by ligand availability alone. Receptor activity is additionally governed by the intracellular itinerary of the receptors themselves: continuous internalisation, endosomal sorting and recycling to the plasma membrane determine how much receptor is available to signal, and for how long. In MFS this itinerary is disturbed. Rather than being routed towards caveolin-1-dependent degradation and signal termination, TGF-β receptors are preferentially internalised through clathrin- and SARA-mediated endocytosis into early endosomes, where SARA recruits SMAD2 and prolongs signalling, while fewer receptors enter normal recycling routes. The result is a feed-forward loop that amplifies SMAD and ERK signalling and contributes to aortic wall weakening.^4^

Vesicular transport is equally central to assembly of the matrix itself. Disorganised fibre architecture is a hallmark of MFS, with fragmented elastic lamellae and disordered collagen networks in the aortic wall.^5^ Collagen folding and fibrillogenesis depend on adequate hydroxylation of proline and lysine residues,^6^ introduced co-translationally in the endoplasmic reticulum (ER) to generate the hydroxyproline [Hyp(P)] and hydroxylysine [Hyp(K)] residues required for triple-helix stability and crosslink formation.^7^

These modifications are catalysed by lysyl oxidases (LOX),^8^ lysyl hydroxylases (PLOD)^9^ and prolyl 3-hydroxylases (P3H),^10^ and their disruption is sufficient to cause aortopathy: LOX loss-of-function variants predispose to TAA and dissection.^11^ Critically, procollagen must transit the ER and Golgi in a tightly regulated manner for these modifications to be completed before secretion, and the Golgi stacking proteins GRASP55 and GRASP65 restrain exocytic transit precisely to safeguard the fidelity of procollagen processing.^12,13^ Both TGF-β receptor availability and collagen maturation therefore depend on the same secretory machinery.

Mixed lineage kinase domain-like protein (MLKL) is best characterised as the terminal effector of necroptosis, a regulated necrotic cell death executed after receptor-interacting protein kinase 3 (RIPK3) phosphorylates its pseudokinase domain (Ser345/Ser347 in mice). Phosphorylated MLKL oligomerizes and translocates to cellular membranes, where it disrupts membrane integrity and causes lytic death.^14–16^ MLKL also operates in membrane and vesicle biology independently of cell killing: it regulates endosomal trafficking and extracellular-vesicle generation through ESCRT and flotillin interactions,^17,18^ is directed to endosomes by site-specific ubiquitination,^19^ participates in antimicrobial defence,^20^ and contributes to tissue remodelling.^21^ In the vasculature, MLKL has been implicated in inflammatory tissue injury including myocardial infarction^22^ and abdominal aortic aneurysm (AAA), where we recently showed that MLKL deficiency limits disease by attenuating SMC necroptosis.^23^ Whether MLKL influences the thoracic aorta, where aneurysm formation is driven by matrix dysfunction rather than inflammation, has not been examined.

Here we asked whether MLKL contributes to aortic wall homeostasis in MFS and, if so, whether that contribution requires necroptotic execution. We used mice carrying a phosphorylation-dead MLKL^S345A/S347A^ (*Mlkl^AA^*) allele, in which MLKL protein is expressed at normal levels but cannot be activated by RIPK3, crossed with the *Fbn1^C1039G/+^* mouse model of MFS. Aortic structure and function, aortic proteome, collagen site-resolved hydroxylation, cell death, and TGFβR1 trafficking in aortic SMCs were assessed.

## Results

### Phosphorylation-dead MLKL causes isolated aortic root dilation with a secretory-pathway proteomic signature

High-frequency ultrasound revealed a modest but significant increase in aortic bulb (root) diameter in 12-week-old *Mlkl^AA^* animals compared with WT littermates, whereas the aortic arch diameter was not significantly enlarged and pulse-propagation velocity (PPV) as a marker for vascular stiffness was unchanged (Fig. 1a-c). The observed phenotype was age-dependent, with no changes in 6-week-old animals (Supplementary Fig. 1a) but with significant increases in bulb and arch diameter in 18-week-old *Mlkl^AA^* as compared to WT mice (Supplementary Fig. 1b). To identify the underlying molecular processes, we performed label-free proteomics on ascending and aortic arch tissue. *Mlkl^AA^*aortas showed a discrete proteomic shift as compared to WT (29 proteins down-regulated, 32 up-regulated; Fig. 1d). Gene-ontology analysis pointed towards overall changes in the secretory and vesicular-trafficking machinery in *Mlkl^AA^*aortas. In particular, down-regulated proteins were enriched for vesicle-mediated transport, regulation of exocytosis, intracellular transport and regulation of SMC migration (Fig. 1e), and included the COPII coat component SAR1B (log_₂_FC = −2.19, *p* = 0.012), the clathrin-adaptor subunit AP1B1 (log_₂_FC = −1.40, *p* = 0.031) and protein kinase C-α (PRKCA; log_₂_FC = −1.25, *p* = 0.001). Up-regulated proteins were enriched for lipid, carboxylic-acid and glycerolipid metabolic processes (Fig. 1f), including fatty-acid synthase (FASN; log_2_FC = +1.22, *p* = 0.021), fatty-acid-binding protein 4 (FABP4; log_₂_FC = +1.53, *p* = 0.005) and the lipogenic regulator THRSP (log_₂_FC = +3.66, *p* = 0.0009).

**Figure 1.**
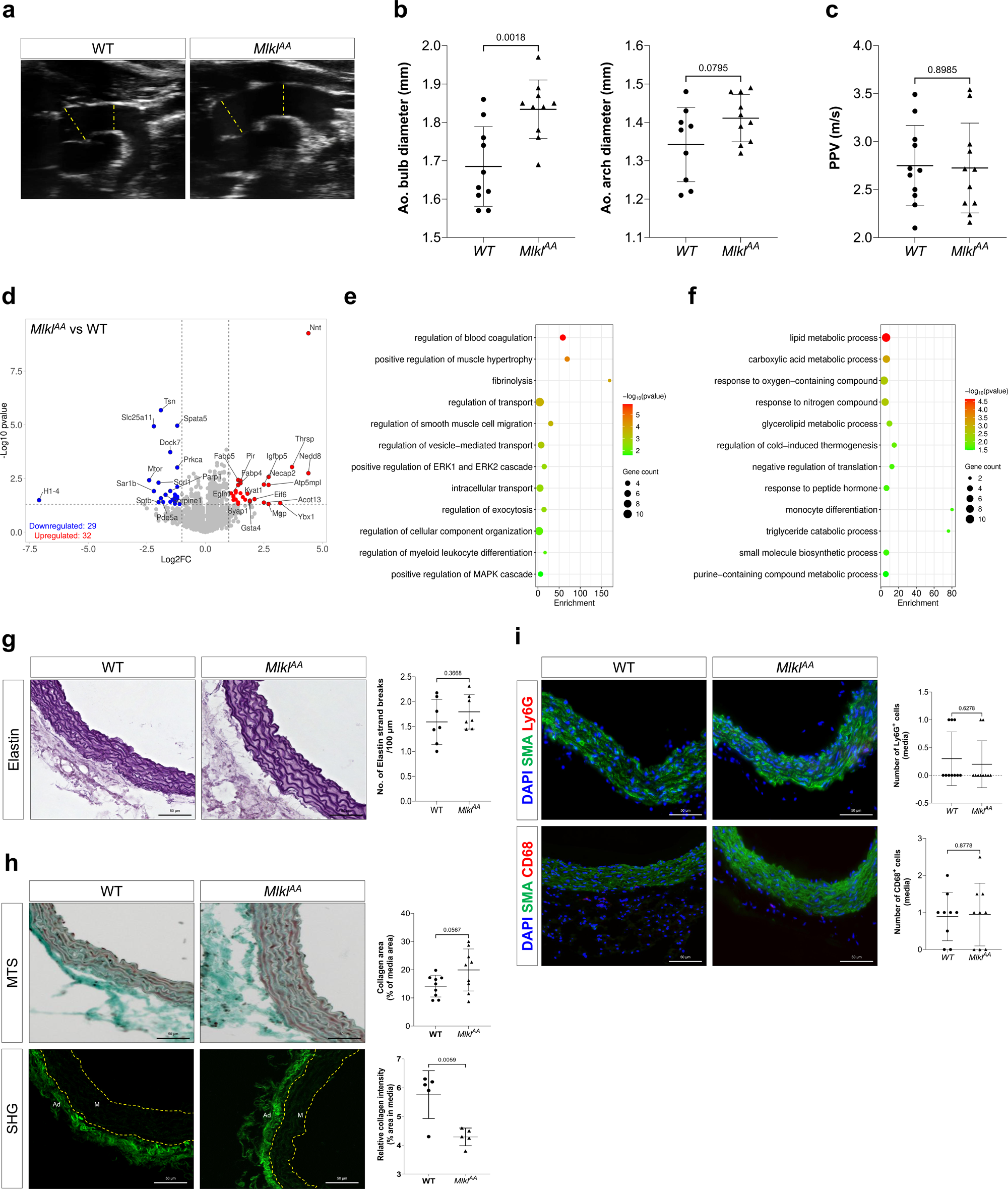
Phosphorylation-dead MLKL causes isolated aortic root dilation with a discrete secretory-pathway proteomic signature. All data from 12-week-old male mice; WT versus *Mlkl* unless stated. (a) Representative B-mode ultrasound images of the thoracic aorta; the dashed yellow line indicates the location of the bulb and arch measurement. (b) Quantification of aortic bulb (left) and arch (right) diameter. (c) Pulse-propagation velocity (PPV). (d) Volcano plot of the bulk aortic proteome depicting differentially abundant proteins in aortic tissue [thresholds: log_₂_FC ≥ 1, *p* < 0.05]. (e and f) Gene Ontology (GO) term enrichment analysis of proteins significantly down-regulated (e) and up-regulated (f) in aortic tissue (dot colour, −log_₁₀_(p); dot size, gene count). (g) Representative elastin staining and quantification of elastic-strand breaks. (h) Representative Masson’s trichrome staining (*n* = 10) and second-harmonic generation (SHG) imaging (*n* = 5) of collagen architecture in the aortic wall. (i) Representative immunofluorescence staining for DAPI (blue), α-SMA (green) and Ly6G (red; top) or CD68 (red; bottom). Data are expressed as mean ± SD. Statistical significance was determined by two-tailed unpaired Student’s t-test (b, c, g, h, and i); *n* = 10 (b, c, g, and i); *n* = 5 (d-f and h).

We next asked whether this proteomic signature was accompanied by altered structural or inflammatory aortic remodelling. In *Mlkl^AA^* animals, elastin architecture was preserved, with no significant increase in elastin-fibre breaks (Fig. 1g). Masson’s trichrome staining, which reflects total collagen abundance, showed only a non-significant trend toward increased medial collagen content compared with WT mice. In contrast, second-harmonic generation (SHG) imaging, which preferentially detects the highly ordered triple-helical structure of fibrillar collagen, revealed a significant disruption of medial collagen fibre organisation in *Mlkl^AA^* aortas, indicating that collagen architecture and structural organisation, rather than overall collagen abundance, were primarily altered (Fig. 1h). Ly6G^+^ neutrophils were essentially absent in both genotypes, while immunofluorescence for CD68^+^ macrophages showed no significant difference between *Mlkl^AA^*and WT (Fig. 1i), indicating that the *Mlkl^AA^ phenotype developed without a detectable inflammatory response*.

### Phosphorylation-dead MLKL aggravates Marfan-associated thoracic aortic dilation

Given the baseline phenotype of *Mlkl^AA^* mice, we next asked whether loss of MLKL activation also impacts established aortic disease, such as MFS. To this end, we crossed *Mlkl^AA^* mice with heterozygous fibrillin-1 mutant (*Fbn1^C1039G/+^*) MFS animals to generate double-mutant *Mlkl^AA^*/*Fbn1^C1039G/+^* mice (DKO^AA^). At 12 weeks, DKO^AA^ mice showed pronounced aortic bulb and aortic arch dilation relative to MFS littermates, and a reduction in PPV (Fig. 2a-c). This aggravation was already present at 6 weeks and persisted at 18 weeks (Supplementary Fig. 1c,d).

**Figure 2.**
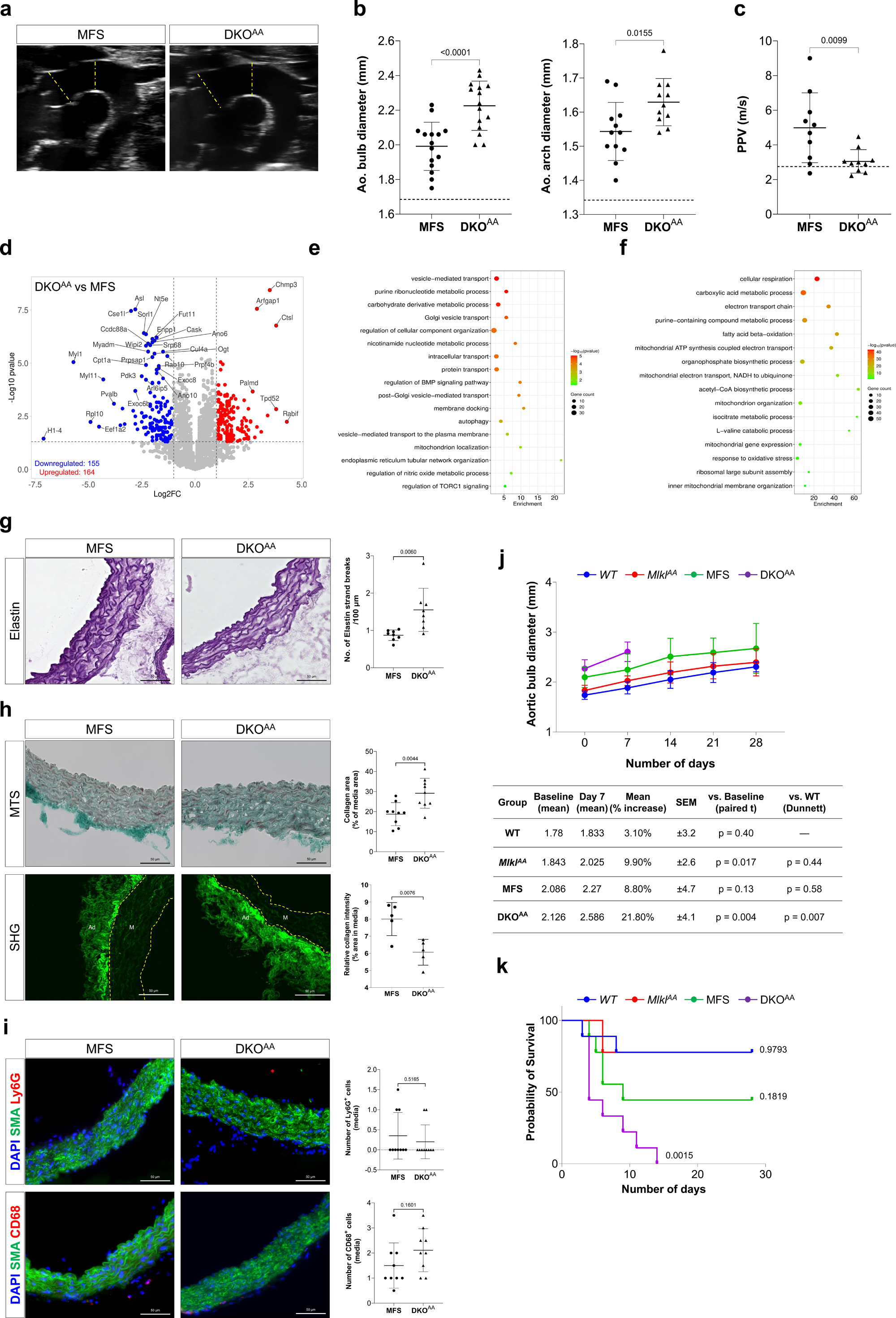
Phosphorylation-dead MLKL aggravates Marfan aortopathy and precipitates lethal aortic disease. All data from 12-week-old male mice; MFS versus DKO^AA^ unless stated. (a) Representative B-mode ultrasound images of the thoracic aorta; the dashed yellow line indicates the location of the bulb and arch measurement. (b) Quantification of aortic bulb (left) and arch (right) diameter. (c) Pulse-propagation velocity (PPV). (d) Volcano plot of the bulk aortic proteome depicting differentially abundant proteins in aortic tissue [thresholds: log_₂_FC ≥ 1, *p* < 0.05]. (e and f) Gene Ontology (GO) term enrichment analysis of proteins significantly down-regulated (e) and up-regulated (f) in aortic tissue (dot colour, −log_₁₀_(p); dot size, gene count). (g) Representative elastin staining and quantification of elastic-strand breaks. (h) Representative Masson’s trichrome staining (*n* = 10) and second-harmonic generation (SHG) imaging (*n* = 5) of collagen architecture in the aortic wall. (i) Representative immunofluorescence staining for DAPI (blue), α-SMA (green) and Ly6G (red; top) or CD68 (red; bottom). (j) Aortic bulb diameter over 28 days of angiotensin II (AngII) infusion in WT, *Mlkl^AA^*, MFS and DKO^AA^ mice (n = 10), with the accompanying table of baseline and day-7 diameters, mean % increase ± SEM, paired t-test versus baseline, and Dunnett’s test versus WT. (k) Kaplan–Meier survival curves depicting the probability of survival in WT, *Mlkl^AA^*, MFS and DKO mice following AngII minipump implantation over a 28-day period. Data are expressed as mean ± SD. Statistical significance was determined by two-tailed unpaired Student’s t-test (b, c, g, h, and i); and by Log-rank (Mantel-Cox) test (k); *n* = 10 (b, c, g, i, j and k); *n* = 5 (d-f and h).

Importantly, ultrasound analysis of a 12-week-old independent *Mlkl^−/−^*cohort reproduced this phenotype. *Mlkl^−/−^* mice exhibited increased thoracic aortic diameter, while MFS mice crossed with *Mlkl^−/−^*mice (DKO^−/−^) recapitulated the aortic dilation along with decreased PPV observed in DKO^AA^ mice (Supplementary Fig. 1e,f). The concordant phenotypes of complete MLKL deficiency and phosphorylation-deficient MLKL indicate that the aortic phenotype results from loss of MLKL function rather than a neomorphic effect of the *Mlkl^AA^* allele.

Proteomic profiling revealed more extensive remodelling in DKO^AA^ aortas than in either *Mlkl^AA^* or MFS aortas (Supplementary Fig. 2), with 155 down-regulated and 164 up-regulated proteins (Fig. 2d). Down-regulated proteins were strongly enriched for vesicle-mediated transport, Golgi vesicle transport, protein transport, post-Golgi vesicle-mediated transport, endoplasmic reticulum tubular network organisation and autophagy (Fig. 2e), reinforcing the secretory-pathway signature seen in *Mlkl^AA^* aortas, and included the endosomal sorting receptor SORL1 (log_₂_FC = −2.40, *p* < 0.0001), the autophagy regulator WIPI2 (log_₂_FC = −2.25, *p* < 0.0001), the exocyst complex component EXOC8 (log_₂_FC = −1.68, *p* < 0.0001), the post-Golgi vesicle GTPase RAB10 (log_₂_FC = −1.28, *p* < 0.0001) and the COPII coat component SAR1B (log_₂_FC = −1.95, *p* = 0.013). By contrast, up-regulated proteins were enriched for cellular respiration, the electron-transport chain, fatty-acid β-oxidation, mitochondrial organisation, mitochondrial gene expression and the response to oxidative stress (Fig. 2f), and included the succinate dehydrogenase complex II SDHB (log_₂_FC = +1.58, *p* = 0.0006), the β-oxidation enzyme HADHA (log_₂_FC = +1.72, *p* = 0.002), the cristae-organising protein CHCHD3 (log_₂_FC = +1.12, *p* = 0.0002) and peroxiredoxin-5 (PRDX5; log_₂_FC = +1.01, *p* = 0.0001).

DKO^AA^ aortas exhibited altered structural remodelling, with a significant increase in elastin-fibre breaks compared to MFS aortas (Fig. 2g), and a significant increase in total medial collagen accompanied by a significant loss of medial collagen fibre organisation as indicated by SHG imaging (Fig. 2h). Ly6G^+^ neutrophils were essentially absent from the media, while CD68^+^ macrophage numbers remained comparable between DKO^AA^ and MFS aortas (Fig. 2i); flow-cytometric quantification of aortic leukocytes likewise showed unchanged proportions of Ly6G^+^ and CD68^+^ cells between all genotypes (Supplementary Fig. 2g).

To determine whether the structural alterations in DKO^AA^ aortas carry functional consequences, we subjected WT, *Mlkl^AA^*, MFS and DKO^AA^ mice to a high dose of angiotensin II (AngII) using an accelerated model of thoracic aortic rupture (Supplementary Fig. 2h).^24^ Aortic bulb diameter increased progressively and most severely in DKO^AA^ mice, with 21.8 ± 4.1% above baseline by day 7 (Fig. 2j). All DKO^AA^ mice died by approximately day 14 from aortic rupture, whereas survival of MFS mice did not differ significantly from WT over the same period (Fig. 2k). Thus, under haemodynamic stress, loss of MLKL phosphorylation markedly aggravates MFS, with clinically relevant consequences for survival.

### MLKL is phosphorylated in the MFS aortic wall without execution of lytic cell death

The aggravation of Marfan aortopathy by a phosphorylation-dead MLKL allele raised the question of whether MLKL is activated in the aortic wall at all and, if so, whether that activation proceeds to necroptotic execution. We therefore quantified MLKL phosphorylated at Ser345 (MLKL^S345^), the RIPK3-dependent modification that licenses necroptosis, across all four genotypes. MLKL^S345^ was detectable in the WT aortic media and was significantly increased in MFS aortas (Fig. 3a).

**Figure 3.**
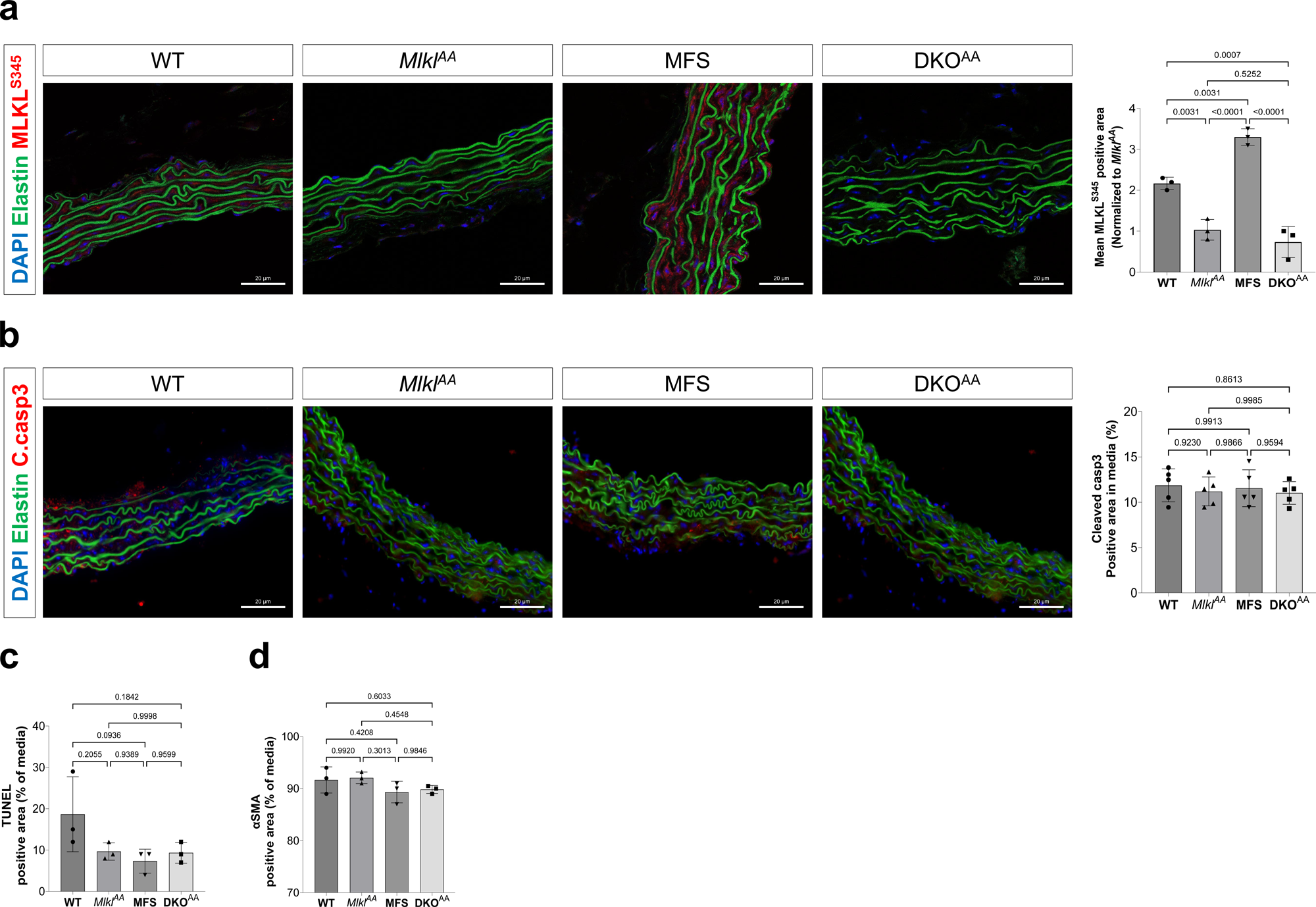
MLKL is phosphorylated in the Marfan aortic wall without execution of cell death. All data from 12-week-old male mice. (a) Left, representative immunofluorescence for MLKL^S345^ (red), elastin (green) and DAPI (blue) in WT, *Mlkl^AA^*, MFS and DKO^AA^ aortas; scale bars, 20 µm. Right, quantification of MLKL^S345^-positive area (% of tissue) and mean MLKL^S345^ intensity. *Mlkl^AA^*and DKO^AA^, which cannot generate the Ser345 phospho-epitope, serve as genetic specificity controls. (b) Left, representative immunofluorescence for cleaved caspase-3 (red), elastin (green) and DAPI (blue); scale bars, 20 µm. Right, cleaved caspase-3-positive medial area (%). (c) TUNEL-positive medial area (%), (d) α-SMA-positive medial area (%). Data are mean ± SD; one-way ANOVA with Tukey’s multiple comparisons test; *n* = 3 (a, c, d), *n* = 5 (b).

We next asked whether this activation was accompanied by cell death. Cleaved caspase-3-positive medial area was comparable across all four genotypes (Fig. 3b), as was the TUNEL-positive medial area (Fig. 3c). Consistent with the absence of cell loss, the α-SMA-positive fraction of the medial area was unchanged across genotypes (Fig. 3d), indicating preserved SMC content.

### Phosphorylation-dead MLKL impairs collagen fibril architecture and proline-lysine hydroxylation in Marfan animals

The secretory-pathway proteomic signature of *Mlkl^AA^*and DKO^AA^ aortas and the altered collagen structure in DKO^AA^ aortas led us to examine SMC collagen ultrastructure and its post-translational modifications. Transmission electron microscopy (TEM) of the aortic tissue across the aortic wall showed progressive disruption of elastic lamellar organisation, SMC morphological changes, ECM disorganisation, and disrupted collagen fibre assembly, most pronounced in DKO^AA^ aortae (Supplementary Fig. 3). Fibril-diameter distributions, tightly clustered in WT, became increasingly heterogeneous in *Mlkl^AA^*, MFS and DKO*^AA^* aortas, with DKO^AA^ showing the most disorganised and small-diameter fibrils (Fig. 4a). In longitudinal sections, collagen D-periodicity, the characteristic ∼67-nm banding pattern of mature fibrils and a measure of fibril structural integrity, was likewise disrupted (Fig. 4b). Because collagen-fibril diameter and D-spacing depend on the post-translational hydroxylation of proline and lysine residues, we performed site-resolved hydroxy-proteomics on aortic tissue.^25^ DKO^AA^ aortas exhibited a broad reduction in prolyl hydroxylation and, to a lesser extent, lysyl hydroxylation across fibrillar collagens, including COL1A1, COL1A2, COL3A1, COL5A1 and COL5A2, compared with WT (Fig. 4c). Among the most significantly affected sites were proline residues in COL1A1 (P761, P767 and P728), COL1A2 (P858, P864, P876 and P882) and COL5A2 (P1196 and P1197), as well as lysine residues in COL1A1 (K731 and K770), COL1A2 (K1014 and K1020) and COL5A1 (K795) (Fig. 4d). This reduction in hydroxylation extended beyond fibrillar collagens to additional ECM, cytoskeletal and TGF-β-regulatory proteins, including APOA1, MCPT8, MYH11, LTBP4, SOD3 and ZNF12 (Supplementary Fig. 4). Among these, LTBP4 controls TGF-β bioavailability in the matrix, while MYH11 and SOD3 act on SMC contractility and oxidative matrix damage. Interestingly, comparison of DKO^AA^ vs MFS aortas revealed a focused set of differential hydroxylation changes (Fig. 4e,f). These sites included COL1A1 (K731, K770, P1153 and P1154) and COL18A1 (P1144 and K1475). Thus, MLKL phosphodeficiency selectively amplifies MFS-associated defects in collagen hydroxylation at specific residues. Reduced hydroxylation also extended to non-collagenous matrix and cytoskeletal proteins, including ACTA2, KANK2, FLNA, MCPT8, HSPA8 and MYH11 (Supplementary Fig. 5).

**Figure 4.**
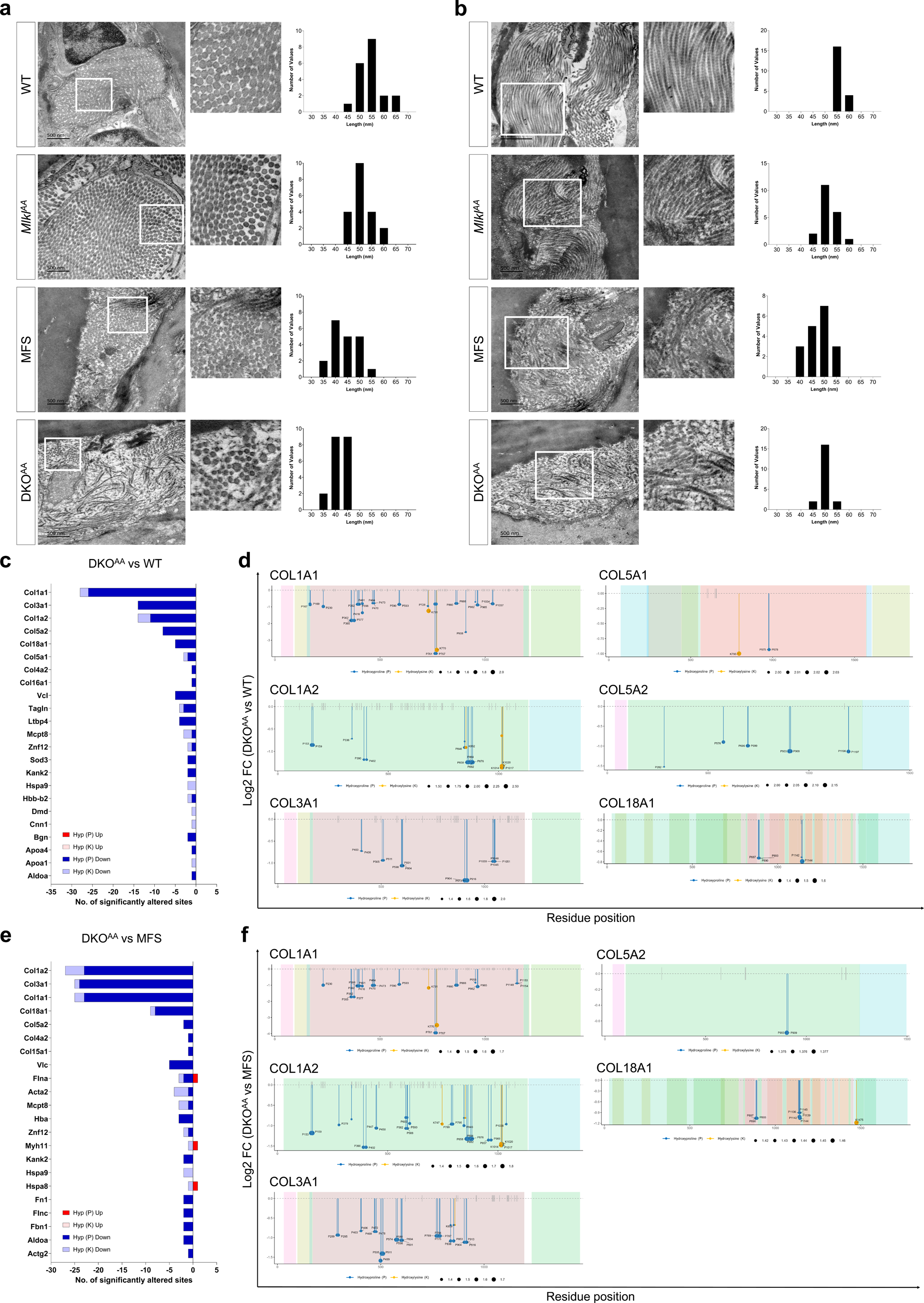
Loss of MLKL activation impairs collagen-fibril architecture and prolyl/lysyl-hydroxylation. (a) Representative TEM of medial collagen in cross-section (fibril diameter), with magnified insets and per-genotype frequency histograms of fibril diameter. (b) Representative TEM of medial collagen in longitudinal section (D-periodicity) with frequency histograms of collagen D-periodicity. (c) Significantly altered prolyl/lysyl hydroxylation sites in DKO^AA^ versus WT aorta; bars coloured by modification and direction (hydroxyproline (P) up/down; hydroxylysine (K) up/down). (d) Lollipop plots mapping significantly altered hydroxylation residues along COL1A1, COL1A2, COL3A1, COL5A1, COL5A2, and COL18A1 in DKO^AA^ versus WT aorta. (e) Significantly altered prolyl/lysyl hydroxylation sites in DKO^AA^ versus MFS aorta. (f) Lollipop plots mapping significantly altered hydroxylation residues along COL1A1 and COL18A1 in DKO^AA^ versus MFS aorta. *n* = 1 (a, b); *n* = 5 (c-f).

Although collagen fibril ultrastructure was altered in *Mlkl^AA^*and MFS aortas (Fig. 4a,b), no significant differences were detected in either *Mlkl^AA^* or MFS versus WT (Supplementary Fig. 6).

### MLKL influences collagen-modifying enzymes and extracellular-matrix programmes in SMCs

To determine whether the observed ECM phenotype is SMC-derived, we generated mouse aortic SMCs stably expressing an *Mlkl*-targeting shRNA (Mlkl-shRNA) or a scrambled control (Scr-shRNA). Mlkl-shRNA reduced *Mlkl* transcript levels by approximately 75-80% (Fig. 5a). MLKL depletion was associated with dysregulation of enzymes involved in collagen hydroxylation and crosslinking. Among the prolyl/lysyl hydroxylases, *Plod1* was down-regulated, whereas *Plod2* was strongly up-regulated (Fig. 5b). Within the 3-hydroxylase family, *P3h2* was up-regulated and *P3h4* down-regulated, while *P3h1* and *P3h3* remained unchanged (Fig. 5c). The lysyl oxidase family was broadly repressed, including *Lox*, *Loxl1*, *Loxl2* and *Loxl4*, whereas *Loxl3* showed an upward trend (Fig. 5d).

**Figure 5.**
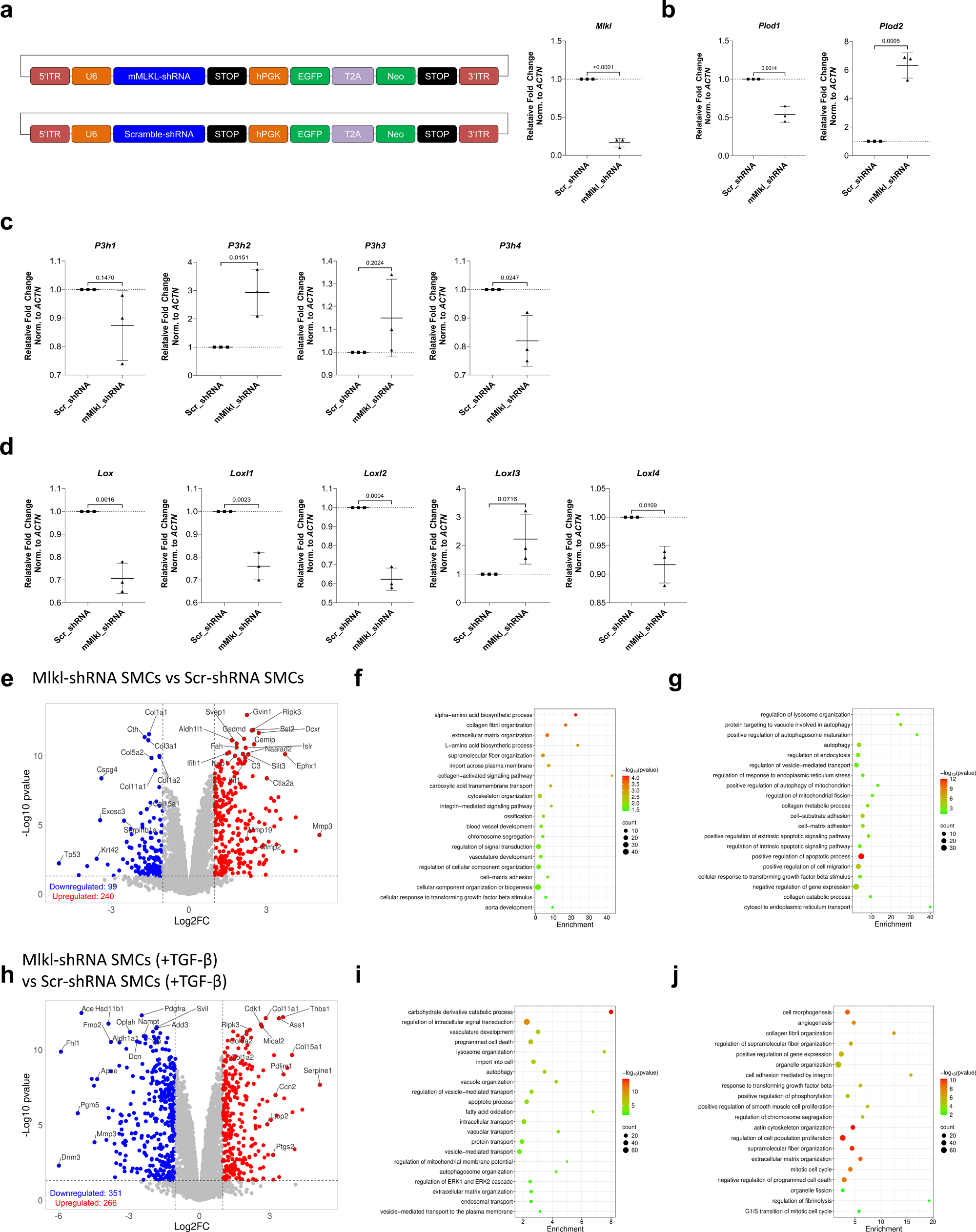
MLKL depletion in smooth muscle cells dysregulates collagen-modifying enzymes and alters TGF-β-driven ECM remodelling. (a) A vector schematic of the Mlkl-shRNA and scrambled-shRNA constructs (left) and *Mlkl* transcript levels represented as relative fold-change, normalised to β-actin (right). (b) Relative fold-change in *Plod1* (left) and *Plod2* (right), normalised to β-actin. (c) Relative fold-change in *P3h1*, *P3h2*, *P3h3* and *P3h4*, normalised to β-actin. (d) Relative fold-change in *Lox*, *Loxl1*, *Loxl2*, *Loxl3* and *Loxl4*, normalised to β-actin. (e) Volcano plot depicting differentially abundant proteins Mlkl-shRNA SMCs versus Scr-shRNA SMCs. (f and g) Gene Ontology (GO) term enrichment analysis of proteins significantly down-regulated (f) and up-regulated (g) in Mlkl-shRNA SMCs versus Scr-shRNA SMCs (dot colour, −log_₁₀_(p); dot size, gene count). (h) Volcano plot depicting differentially abundant proteins in TGF-β treated Mlkl-shRNA SMCs versus Scr-shRNA SMCs. (i and j) Gene Ontology (GO) term enrichment analysis of proteins significantly down-regulated (i) and up-regulated (j) in TGF-β treated Mlkl-shRNA SMCs versus Scr-shRNA SMCs, (dot colour, −log_₁₀_(p); dot size, gene count). Data are expressed as mean ± SD. Statistical significance was determined by two-tailed unpaired Student’s t-test (a, b, c, and d); *n* = 3 (a-j).

Global proteomic profiling of Mlkl-shRNA SMCs vs Scr-shRNA SMCs identified 99 down-regulated proteins (Fig. 5e) including fibrillar collagens COL3A1 (log_2_FC = −1.56, *p* < 0.0001), COL1A1 (log_2_FC = −1.53, *p* < 0.0001), COL5A2 (log_2_FC = −1.44, *p* < 0.0001) and COL11A1 (log_₂_FC = −1.29, *p* < 0.0001), together with the proteoglycan CSPG4 (log_2_FC = −3.37, *p* < 0.0001) were enriched for GO terms such as collagen-fibril organisation, ECM organisation, supramolecular-fibre organisation, collagen-activated signalling and the response to TGF-β (Fig. 5f). In contrast, 240 up-regulated proteins, including lysosomal proteases cathepsin D (CTSD; log_2_FC = +1.15, *p* < 0.0001) and cathepsin H (CTSH; log_2_FC = +1.91, *p* < 0.001), the sphingolipid-activator prosaposin (PSAP; log_2_FC = +1.59, *p* < 0.0001), the collagenase MMP3 (log_2_FC = +5.05, *p* < 0.001) and the adhesion protein SVEP1 (log_2_FC = +1.85, *p* < 0.0001) were enriched for GO terms such as autophagy, lysosome organisation, endoplasmic-reticulum-stress responses, mitochondrial fission and collagen catabolism (Fig. 5g).

Hinting at a potential shift in SMC phenotype and ECM turnover, Mlkl-shRNA SMCs also showed increased mRNA expression of *Acta2* and *Tnfrsf11b* and decreased *Klf4*, together with up-regulation of *Mmp2* and *Mmp13* and down-regulation of *Mmp9* (Supplementary Fig. 7).

Because TGF-β is a key modulator of aortic ECM remodelling and a driver of MFS pathophysiology, we asked whether MLKL shapes SMC responses to TGF-β. Principal component analysis of all four conditions separated samples along the first two principal components (PC1, 38.5%; PC2, 19.1% of variance; Supplementary Fig. 8a). Comparing each genotype with and without TGF-β showed that the response to the ligand itself depended on MLKL: TGF-β altered the abundance of 625 proteins in Scr-shRNA SMCs but only 221 in Mlkl-shRNA SMCs, with the induction arm particularly affected (325 versus 53 up-regulated proteins). The greater divergence between genotypes under TGF-β treatment (618 proteins) than at baseline (339 proteins) therefore reflects a diminished, rather than an exaggerated, response to TGF-β in MLKL-depleted cells (Supplementary Fig. 8b). Global proteomic profiling showed a markedly altered response to TGF-β treatment between Mlkl-shRNA SMCs and Scr-shRNA SMCs with 351 down-regulated proteins (Fig. 5h) including angiotensin-converting enzyme (ACE; log_2_FC = −5.02, *p* < 0.0001), PDGFRA (log_2_FC = −2.45, *p* < 0.0001), decorin (DCN; log_2_FC = −2.65, *p* < 0.0001), apolipoprotein E (APOE; log_2_FC = −4.35, *p* < 0.0001) and FHL1 (log_2_FC = −5.89, *p* < 0.0001); were enriched for GO terms such as intracellular signal transduction, vesicle-mediated transport, vacuolar organisation and transport, endosomal transport and autophagy (Fig. 5i). In contrast, 267 up-regulated proteins, including SERPINE1/PAI-1 (log_2_FC = +5.14, *p* < 0.0001), thrombospondin-1 (THBS1; log_2_FC = +3.57, *p* < 0.0001), the matricellular protein CCN2/CTGF (log_2_FC = +3.24, *p* < 0.0001), COL11A1 (log_2_FC = +2.83, *p* < 0.0001) and the proliferation marker CDK1 (log_2_FC = +2.62, *p* < 0.0001); were enriched for GO terms such as collagen-fibril organisation, smooth-muscle-cell proliferation, integrin-mediated adhesion and the response to TGF-β (Fig. 5j). Notably, several collagens and matricellular proteins that were down-regulated by MLKL depletion in untreated cells, including COL11A1 and CCN2, were instead elevated after TGF-β treatment, indicating that loss of MLKL redirects rather than uniformly suppresses the TGF-β-driven matrix programming in SMCs.

### MLKL activity is required for TGF-β-receptor trafficking and secretory-pathway organisation in SMCs

Given the overlap between the proteomic signatures in *DKO^AA^*aortic tissue and Mlkl-depleted SMCs, pointing to altered TGF-β signalling and receptor trafficking, we first investigated TGFβR1 protein abundance. Immunofluorescence staining revealed a significant increase in TGFβR1 abundance in DKO^AA^, whereas *Mlkl^AA^* and MFS did not differ significantly from WT aortas. TGFβR1 abundance was also significantly higher in DKO^AA^ aortas than in MFS aortas (Fig. 6a).

**Figure 6.**
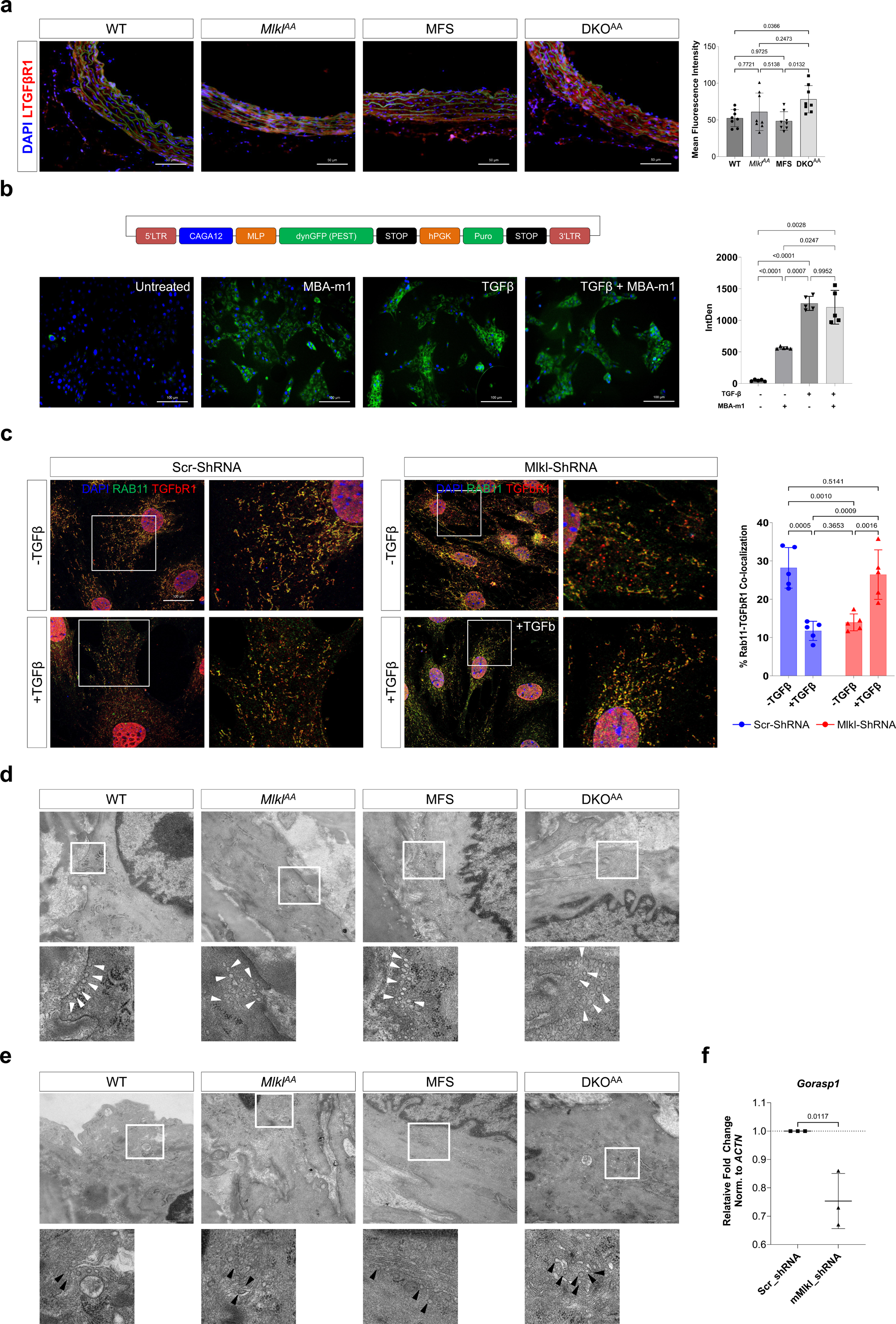
MLKL activity is required for TGF-β-receptor trafficking and secretory-pathway organisation in smooth muscle cells. (a) Left, representative immunofluorescence images for TGFβR1 (red) and DAPI (blue) and right, TGFβR1 quantification in WT, *Mlkl^AA^*, MFS and DKO aortas. (b) Top, a vector schematic of the CAGA_12_-driven destabilised-GFP TGF-β/SMAD reporter (pLV-CAGA-dynGFP). Bottom, representative fluorescence images of reporter in mSMCs, with integrated-density quantification (right). (c) Left, representative immunofluorescence images for TGFβR1 (red), RAB11 (green) and DAPI (blue) and right, RAB11-TGFβR1 co-localisation quantification; in Scr-shRNA and Mlkl-shRNA mSMCs ± TGF-β. (d) Representative TEM images of medial SMCs showing secretory vesicles (white arrowheads) in WT, *Mlkl^AA^*, MFS and DKO aortas. (e) Representative TEM images of medial SMCs showing Golgi-associated budding vesicles (black arrowheads) in WT, *Mlkl^AA^*, MFS and DKO aortas. (f) Relative fold-change in *Gorasp1*, normalised to β-actin in Mlkl-shRNA SMCs versus Scr-shRNA SMCs. Data are expressed as mean ± SD. Statistical significance was determined by one-way ANOVA with Tukey’s multiple comparisons test (a, b, and c); *n* = 8 (a); *n* = 5 (b, c); *n* = 1 (d, e); *n* = 3 (f).

To test whether MLKL activity influences TGF-β signalling, we used a CAGA_₁₂_-driven GFP reporter of SMAD2/3/4 transcriptional activity in murine SMCs. Pharmacological MLKL inhibition with MBA-m1 significantly increased TGF-β signalling activity relative to untreated cells. TGF-β was included as a positive control and produced the strongest reporter activity, which MBA-m1 did not further increase (Fig. 6b).

TGF-β receptors are continuously internalised and recycled to the plasma membrane through RAB11-positive endosomes, a trafficking route that helps control TGF-β signalling by regulating receptor availability at the cell surface.^26,27^ To test whether heightened TGFβR1 surface expression and activity are mediated by impaired trafficking, we assessed the co-localisation with RAB11. In control Scr-shRNA SMCs, TGF-β treatment reduced RAB11-TGFβR1 co-localisation, consistent with ligand-induced redistribution of TGFβR1 away from the recycling compartment. In contrast, Mlkl-shRNA SMCs showed the opposite response: RAB11–TGFβR1 co-localisation was already reduced at baseline (≈14%) but increased following TGF-β treatment (≈27%), reaching levels comparable to those in untreated control cells (Fig. 6c). Thus, MLKL depletion reverses the normal TGF-β-induced redistribution of TGFβR1, suggesting that MLKL contributes to the regulation of TGF-β signalling through the recycling-endosome compartment.

Consistent with the proteomic signature of altered vesicle transport in DKO^AA^ aortas, TEM showed a progressive accumulation of secretory vesicles in aortic SMCs of all three genotypes relative to WT, most pronounced in DKO^AA^ (Fig. 6d) and displaying more Golgi-associated budding vesicles (Fig. 6e), indicating the most pronounced disruption of the vesicle transport apparatus.

Next, we examined the Golgi-stacking protein GRASP65, encoded by *Gorasp1*, as an additional marker of disturbed vesicular transport.^28^ Indeed, Mlkl-shRNA SMCs showed reduced *Gorasp1* mRNA levels (Fig. 6f), consistent with the accelerated, disorganised vesicular trafficking observed in MLKL-phosphodeficient and MFS aortas.

The alterations in the Golgi apparatus extended to the ER, as DKO^AA^ aortas exhibited shortened ER profiles and visibly fewer mitochondria-ER contact sites (MERCs), membrane microdomains that mediate metabolite transfer between the ER and mitochondria. Mitochondria also appeared smaller in both MFS and DKO^AA^ aortas (Supplementary Fig. 9). These findings link the defect in secretory trafficking to the mitochondrial reprogramming observed in the DKO^AA^ proteome.

These data indicate that MLKL activity is required for the organisation of the SMC secretory pathway and for TGFβR1 trafficking. Its loss is associated with dysregulated basal TGF-β signalling, possibly via an altered RAB11-dependent TGFβR1-recycling response, and accumulation of secretory and Golgi vesicles, all occurring in the absence of lytic cell death function, and thus consistent with a non-lytic, homeostatic function of MLKL in SMCs.

## Discussion

MLKL mediates necroptosis following RIPK3-dependent phosphorylation of its pseudokinase domain (S345/S347 in mice). This phosphorylation activates MLKL, enabling its N-terminal four-helix bundle to disrupt cell membranes.^29^ Using mice carrying a phosphorylation-deficient *Mlkl^S345A/S347A^* allele, in which MLKL is expressed normally but cannot be activated by RIPK3, we found that MLKL activation is required for thoracic aortic homeostasis. We found phosphorylated MLKL in the baseline aortic media, which was significantly increased in MFS aortas, while cleaved caspase-3, TUNEL positivity, medial SMC content and leukocyte infiltration remained unchanged, suggesting MLKL activation in the thoracic aortic wall may serve a non-lytic function.

*Mlkl^AA^* mice displayed an enlarged aortic bulb, accompanied by altered collagen organisation within the aortic wall. Total medial collagen content was not significantly increased, whereas SHG imaging revealed a significant loss of fibre organisation, indicating that collagen architecture rather than abundance was primarily affected. These changes occurred without a detectable increase in inflammatory cell content in the aortic wall, despite the well-established link between MLKL and inflammation.^17,23^

Disorganised collagen fibres are a hallmark of MFS and contribute to early vessel rupture in patients,^30^ arising when mutations in FBN1 remove the microfibrillar scaffold that organises collagen deposition. Because D-periodicity is a key determinant of fibril organisation and biomechanical properties, the altered D-periodicity in DKO^AA^ aortas reflects nanoscale disruption of fibril assembly.^31^

Excessive TGF-β activation independently dysregulates collagen folding in vascular SMCs, including impaired prolyl and lysyl hydroxylation.^32,33^ The hydroxylation changes we identified occurred predominantly in collagens,^34^ at or near the residues with established roles in collagen cross-linking and fibril assembly e.g. in COL1A1 near K87 and K930, whose site-specific under-hydroxylation alters cross-linking and reduces fibril diameter without impairing triple-helix formation,^35,36^ and in COL5A1/COL5A2, minor collagens that regulate fibril nucleation and diameter through heterotypic interaction with type I collagen.^37^

TGF-β signalling controls components of the hydroxylation and cross-linking machinery via regulation of LOX and PLOD2,^38^ while P3H1-mediated hydroxylation suppresses TGF-β signalling,^39^ indicating reciprocal regulation. Notably, we found LOX, PLOD, and P3H family members to be dysregulated under MLKL knockdown, linking MLKL to this axis.

MLKL knockdown altered the SMC response to TGF-β in two complementary ways. MLKL inhibition alone activated SMAD-dependent transcription, indicating elevated basal receptor signalling, while the global proteomic response to exogenous TGF-β was markedly reduced. Elevated basal signalling combined with a diminished response to added ligand is consistent with a chronically engaged, desensitised receptor system. TGF-β receptor activation depends not only on ligand-induced dimerization and phosphorylation but on a vesicle-dependent cycle of endocytosis, recycling and return to the membrane,^40^ governed by vesicular transport;^41^ signalling occurs at the plasma membrane and, more efficiently, at early endosomes,^42,43^ a balance that requires fine control.^44^ MLKL interacts with essential components of this cycle and can interrupt, delay or accelerate receptor trafficking.^45^ Our data fit these observations directly: in MLKL-depleted cells, less TGFβR1 resided in the RAB11 recycling compartment at baseline, while TGF-β drove receptor into that compartment rather than away from it. Within this blunted overall response, a subset of fibre-associated collagens and matricellular proteins, including COL11A1 and CCN2, was selectively induced, indicating that MLKL loss redirects rather than uniformly suppresses TGF-β-driven matrix programming.

Disruptions in both Golgi and ER architecture appeared in *Mlkl^AA^*aortas and were more pronounced in DKO^AA^ aortas, accompanied by changes in mitochondrial structure, consistent with the functional coupling between the secretory pathway and mitochondrial homeostasis through ER- and Golgi-derived trafficking and inter-organelle contact sites, such that defects in the ER–Golgi network could indirectly impair mitochondrial integrity.^46^

How MLKL acts within the vesicle compartment is not fully established, but MLKL is known to be associated with flotillins (flotillin-1/2) and participate in flotillin-mediated endocytosis.^47^ In addition, MLKL interacts with the ESCRT machinery that mediates multivesicular-body and extracellular-vesicle biogenesis,^17^ and with the retromer complex (VPS35-VPS29-VPS26), which controls RAB7A activity and the endosomal sorting/recycling system.^48^ Relevant to our model, site-specific ubiquitination directs MLKL to endosomes, where it promotes endosomal trafficking and vesicle release independently of necroptotic signalling, this ubiquitination occurs at sites distinct from the Ser345/Ser347 residues mutated in *Mlkl^AA^*.^19^ Endosomal MLKL trafficking may therefore remain intact in *Mlkl^AA^* and redirect MLKL towards a non-lytic endosomal pathway that supports RAB11-dependent TGFβR1 recycling in the aortic wall. Consistent with this hypothesis, the accumulation of secretory and Golgi-associated vesicles in MLKL-phosphodeficient aortas, together with aggravated aortic dilation, disorganised collagen architecture and reduced survival in DKO^AA^ mice, suggests that preventing MLKL phosphorylation may disrupt a compensatory, potentially protective response.

These findings also account for an apparent discrepancy with our previous report that MLKL deficiency protects AAA by attenuating SMC necroptosis.^23^ Two differences between AAA and MFS are likely relevant. SMCs of the aortic root and ascending aorta derive from cardiac neural crest and second heart field, whereas those of the abdominal aorta arise from splanchnic mesoderm, influencing SMC phenotype and the response to TGF-β.^49^ Furthermore, susceptibility to aortic diseases is SMC lineage dependent.^50^ Second, the inflammatory context differs, AAA is considered a pro-inflammatory condition driven by pronounced leukocyte infiltration, whereas MFS aortas show comparatively low-grade inflammation with substantially less inflammatory cell accumulation.^51^

Together, these findings identify a non-lytic, homeostatic role for MLKL in vascular SMCs, in which MLKL supports vesicle organisation and TGFβR1 trafficking. Loss of this function is associated with impaired RAB11-dependent TGFβR1 recycling, accumulation of secretory and Golgi vesicles, and dysregulated basal TGF-β signalling, without execution of lytic cell death. MLKL may therefore be injurious or protective in the vessel wall depending on aortic segment and inflammatory context. Whether this extends to other TGF-β-driven aortopathies, such as Loeys-Dietz syndrome^52^ and systemic sclerosis^53^ remains to be investigated in future studies.

### Limitations

1. The trafficking mechanism we describe, RAB11-dependent TGFβR1 recycling and Golgi-associated vesicle accumulation, is established in cultured aortic SMCs, and an SMC-specific *Mlkl* deletion would be required to attribute the aortic phenotype unambiguously to SMC-intrinsic MLKL function.
2. All experiments were performed in male mice; given the sex-dimorphism of Marfan aortopathy, our findings cannot be extrapolated to female animals.
3. Our data establish that MLKL phosphorylation is increased in MFS aortas and that preventing it aggravates disease, but they do not identify the upstream signal driving MLKL phosphorylation in the aortic wall, nor the molecular interaction through which phosphorylated MLKL engages the recycling machinery.
4. The cultured SMCs used here were commercially sourced aortic SMCs of unspecified segmental origin; given the lineage differences between thoracic and abdominal aortic SMCs, segment-matched cells would be required to establish that the trafficking phenotype is thoracic-specific.

## Methods

### Animal studies

Animal studies were approved by the local animal care authorities, Landesamt für Natur, Umwelt und Verbraucherschutz Nordrhein-Westfalen (LANUV) (AZ: 2021.A004) and conformed to the guidelines from Directive 2010/63/EU of the European Parliament on the protection of animals used for scientific purposes. All experiments were performed on male mice and littermates were used as controls. All animals were given free access to a standard chow (Altromin 1324 P Best, Altromin GmbH & Co. KG, Lage, Germany) and water; all interventions were performed under isoflurane anaesthesia with buprenorphine analgesia as per the LANUV guidelines. All experimental animals were monitored daily and were given proper medical attention with appropriate care and sacrificed when reaching pre-determined criteria of disease severity. No specific randomisation protocol had been used to select animals. We aimed to analyse at least 10 animals per group in a blinded fashion. Animals were maintained at the animal facility of the University of Cologne under strict SPF housing conditions, at 22 °C (±2 °C) with 55% (±5%) humidity in individually ventilated cages under a 12-h light cycle.

### Mouse strains and genotyping

Marfan mice carrying the *Fbn1^C1039G/+^* allele^54^ (strain #012885, Jackson Laboratory) were crossbred with phosphorylation-dead *Mlkl^S345A/S347A^* (*Mlkl^AA^*) and *Mlkl^−/−^* mice^55^ to generate DKO^AA^ (*Fbn1^C1039G/+^*;*Mlkl^AA^*) and DKO^-/-^ (*Fbn1^C1039G/+^*;*Mlkl^-/-^*) animals. Genotyping was performed by PCR using the following primers: Mlkl_F, 5’-CATCAAGTTAGGCCAGCTCA -3’, Mlkl_R, 5’- TCTGCTGGTTAGCCTCCTTC -3’; MlklAA_KO_F 5’-AAAACACAGAATGCCATCGC -3’, MlklAA_KO_R 5’- GTGGAAGGAGAGAAATGGCCCT -3’; Mlkl_WT_F 5’- GCAAAACACAGAATTCCATCAG -3’, Mlkl_WT_R, 5’-GTGGAAGGAGAGAAATGGCCCT -3’; MFS_F 5’- CTCATCATTTTTGGCCAGTTG -3’, MFS_R 5’- GCACTTGATGCACATTCACA -3’. All mice were backcrossed and maintained on a C57BL/6N background.

### Angiotensin II infusion model

12-week-old male mice were challenged by continuous subcutaneous infusion of angiotensin II (AngII) (A9525, Sigma-Aldrich) dissolved in 2 µL of 0.5 M acetic acid per 1 mL of saline at 3125 ng/kg/min, delivered by osmotic minipumps (model 1002; ALZET), implanted under isoflurane anaesthesia. Pumps were replaced on day 14 to maintain delivery over the 28-day period, and mice were monitored by weekly ultrasound.^56^

### Ultrasound imaging of the thoracic aorta

Aortic ultrasound was performed as previously described by Nemade et al.^23^ In brief, mice were anaesthetized with 2% isoflurane, and the thoracic aorta was imaged in B-mode, M-mode and EKV on a Vevo3100 system with an MX550D (25-55 MHz, center transmit: 40 MHz, axial resolution: 40 μm) linear-array transducer (VisualSonics). The aortic bulb and ascending arch were measured, and absolute luminal diameters (mm) together with pulse-propagation velocity (PPV) were quantified from B-mode/EKV recordings using Vevo software (Version). Ultrasound measurements were acquired at baseline, 7, 14, 21, and 28 days of the AngII-infusion experiment whereas, for the disease progression models, the measurements were acquired at 6, 12, and 18 weeks of age.

### Histology

Masson’s trichrome staining and elastin staining were performed on aortic cryosections as previously described by Nemade et al.^23^ All Images were acquired on a BZ-X800 (Keyence, USA) and analysed using ImageJ software. Image acquisition and quantification were performed separately for each cohort, absolute values are comparable within but not between cohorts, and no statistical comparisons were made across cohorts.

### Immunofluorescence staining

Thoracic aorta was harvested, cleaned and embedded into Tissue-Tek O.C.T. Compound (Sakura Finetek™ 4583). After freezing at -80 °C, the tissue was cut into 10 µm-thick sections using a Leica CM3050 S Cryostat (Leica, Germany), fixed with 4% paraformaldehyde (PFA) for 10 min and permeabilized using 0.1% Triton X-100 for 10 min at room temperature. Sections were blocked with 3% bovine serum albumin (BSA) for 1 h at RT and incubated with the primary antibody at 4 °C overnight (Supplementary Table S1). Subsequently, sections were washed and incubated with species-matched secondary antibodies for 1 h at RT in the dark, washed and mounted with ProLong^TM^ Glass Hard-Set Antifade Mounting Agent (P36980, Thermo Fisher, USA). Images were acquired with a BZ-X800 microscope system (Keyence, USA) and analysed using ImageJ software.

### Proline/lysine hydroxylation proteomics

Hydroxylation (proline/lysine) site tables provided by the CECAD proteomics core facility (Cologne, Germany) were analysed in Perseus (v1.6.15.0). Site-level signals were compared between genotype groups by two-sided Student’s t-test with S0 = 0.2 and permutation-based FDR = 0.05 (250 randomisations); sites passing these thresholds were considered significant. Figures were generated in R (v4.5.1) with ggplot2. Bar plots show, for each protein, the number of residue positions at which hydroxylation was significantly changed with an absolute log_₂_ difference of at least 0.585. Lollipop plots show significant hydroxylation sites along the protein sequence; stem height represents the log_2_ difference, colour indicates the modified residue, and point size represents −log_10_(q). Each site was positioned according to its amino-acid position in the corresponding mouse UniProt sequence, with UniProt domain and region annotations displayed in a schematic map beneath each protein.

### Flow cytometry of aortic leukocytes

Aortas from 12-week-old mice were perfused, enzymatically digested, processed to single-cell suspensions, and stained with antibodies previously described in Nemade et al.^23^ Samples were acquired on a 5-laser Cytek^®^ Aurora and analysed in FlowJo. Immune cell populations were identified as CD45^+^, Zombie UV^-^ and subpopulations were defined as neutrophils (CD11b^+^, CD3^-^, Ly6G^+^); macrophages (CD11b^+^, CD3^-^, F4/80^+^, MHCII^+^).

### In vitro methods Reagents

Mouse primary aortic smooth muscle cells were received from Cell Biologics (C57-6080) and cultured in the complete smooth muscle cell medium (M2268, Cell Biologics) as per the manufacturer’s protocol in a standard cell culture incubator at 5% CO , 37 °C. All other cell culture reagents, such as DMEM, RPMI, TrypLE™ select enzyme, FBS, HBSS buffer, etc., were purchased from Gibco/Thermo Scientific (Waltham, MA, USA). Geneticin (G418) was purchased from InvivoGen (ant-gn-2, InvivoGen).

### Generation of Mlkl knockdown SMCs

Stable Mlkl knockdown was achieved using a piggyBac vector encoding an Mlkl-targeting shRNA (target sequence: 5′-AGATCCAGTTCAACGATATAT-3′) driven by the U6 promoter, together with an EGFP-T2A-Neo selection cassette; a scrambled shRNA in the same backbone served as the control (Scr-shRNA) (VectorBuilder, USA). A stable SMC line was generated by transfecting the shRNA vector along with pRP-mCherry-CAG-hyPBase transposase vector using FuGENE^®^ HD Transfection Reagent (E2311, Promega, USA) as per the manufacturer’s protocol. Cells were selected, 48 h post-transfection, with 800 µg/ml G418 for 6 days, after which the G418 concentration was reduced to 400 µg/ml. Knockdown efficiency was confirmed by qRT-PCR.

### TGF-β reporter assay

TGF-β/SMAD transcriptional activity was measured using the CAGA_₁₂_-driven destabilised-GFP reporter (pLV-CAGA-dynGFP-Neo), kindly provided by Dr. Peter ten Dijke,^57^ transiently transfected into mSMCs with FuGENE^®^ HD (Promega). Cells were treated with TGF-β1 (763104, Biolegend, USA) at 5 ng/ml for 24 h, alone or in combination with the MLKL inhibitor MBA-m1 at 1000 ng/ml and reporter GFP signal was quantified as integrated density in ImageJ.

### RAB11-TGFβR1 co-localisation

The mSMCs (Scr-shRNA or Mlkl-shRNA) were serum-starved for 24 h by culturing in 0.1% FBS media then treated with 5 ng/ml TGF-β1 for 24 h, fixed and co-stained for RAB11 and TGFβR1, and Images were acquired using a STELLARIS 8 STED confocal laser scanning microscope (Leica Microsystems, CECAD). Co-localisation was quantified using the ComDet ImageJ plugin and represented as % co-localisation.

### Quantitative real-time PCR

Total RNA was isolated using RNeasy Plus Mini Kit (74134, QIAGEN, Germany). 500 ng of total RNA was then reverse-transcribed using High-Capacity cDNA Reverse Transcription Kit (4368814, Thermo Fisher, USA) and analysed by QuantStudio 1 Real-Time-PCR-System (Applied Biosystems, USA) using GoTaq^®^ qPCR master mix (A6001, Promega, USA). Transcripts were normalised to Actb and expressed as relative fold change (2^^−ΔΔCt^). Primer sequences for all tested genes are shown in supplementary table S2.

### Statistics

Data are presented as mean ± SD. Normality and equality of variances were assessed by Shapiro–Wilk and Brown-Forsythe tests. Two groups were compared by two-tailed unpaired Student’s t-test; three or more groups by ordinary one-way or two-way ANOVA with Tukey’s correction for multiple comparisons. For AngII survival experiments, the survival was analysed by Log-rank (Mantel-Cox) test. *p* < 0.05 was considered significant. Analyses used GraphPad Prism 10.

TEM was performed on a single animal per genotype. Because individual fibrils, vesicles and organelles measured within one animal do not constitute independent biological replicates, no inferential statistics were applied to ultrastructural measurements. Collagen fibril diameter and D-periodicity, secretory and Golgi-associated vesicle numbers, and mitochondrial and endoplasmic-reticulum dimensions (Figs. 4a,b and 6d,e; Supplementary Figs. 3 and 9) are reported as descriptive distributions or counts pooled from 20 fields per animal, and all genotype comparisons based on these measurements are qualitative.

## Funding

Deutsche Forschungsgemeinschaft (DFG, German Research Foundation)-Grant No. 397484323 - CRC/TRR259; projects A04, A05, A09, B05, B09 and C03 to MM, MA, HW, StB and GS. The Köln Fortune Program and the Walter und Maga Boll Foundation to TR.

## Disclosure

All the authors declared no competing interests.

## Acknowledgments

The authors would like to thank Christina Vosen, Katharina Tinaz, Nadja Klein, and Simon Grimm for their excellent technical support. We acknowledge support from the Cologne Excellence Cluster on Cellular Stress Responses in Aging Associated Diseases (CECAD) Proteomics Core Facility (Dr. Jan-Wilm Lackmann and Dr. Stefan Müller) and from the CECAD Imaging Facility (Dr. Astrid Schauss). We also acknowledge the proficient technical support by Dr. Matthias Mörgelin (Colzyx AB, Lund, Sweden) in TEM analysis.

## Contributions

MM and MA conceived and supervised the study. HN harvested the animals, gathered and analysed data, and wrote the manuscript. HG harvested the animals, gathered and analysed data, performed echocardiography, histology and immunofluorescence staining, proteomics analysis, cell culture, and animal experiments. OZ, SR, MH, HS, KML, BA, DZ, FH, PA, DM, PJ, and FN performed echocardiography, histology and immunofluorescence staining, proteomics analysis, cell culture and animal experiments. PS, FRP and HW performed FACS and analysed the data. JJ and GS performed hydroxylation proteome analysis. UR and AGS supplied the MLKL inhibitor, MBA-m1. TR, SB, HW, MA, and MM provided experimental support with mouse lines, surgeries, and ethics applications. All authors participated in data analysis, critically reviewed the manuscript, and gave final approval.

## Notes

### Competing Interest Statement

The authors have declared no competing interest.

