## Supplementary Figure 1-9 for "A cell death-independent role for MLKL in TGF-β receptor recycling in Marfan syndrome"

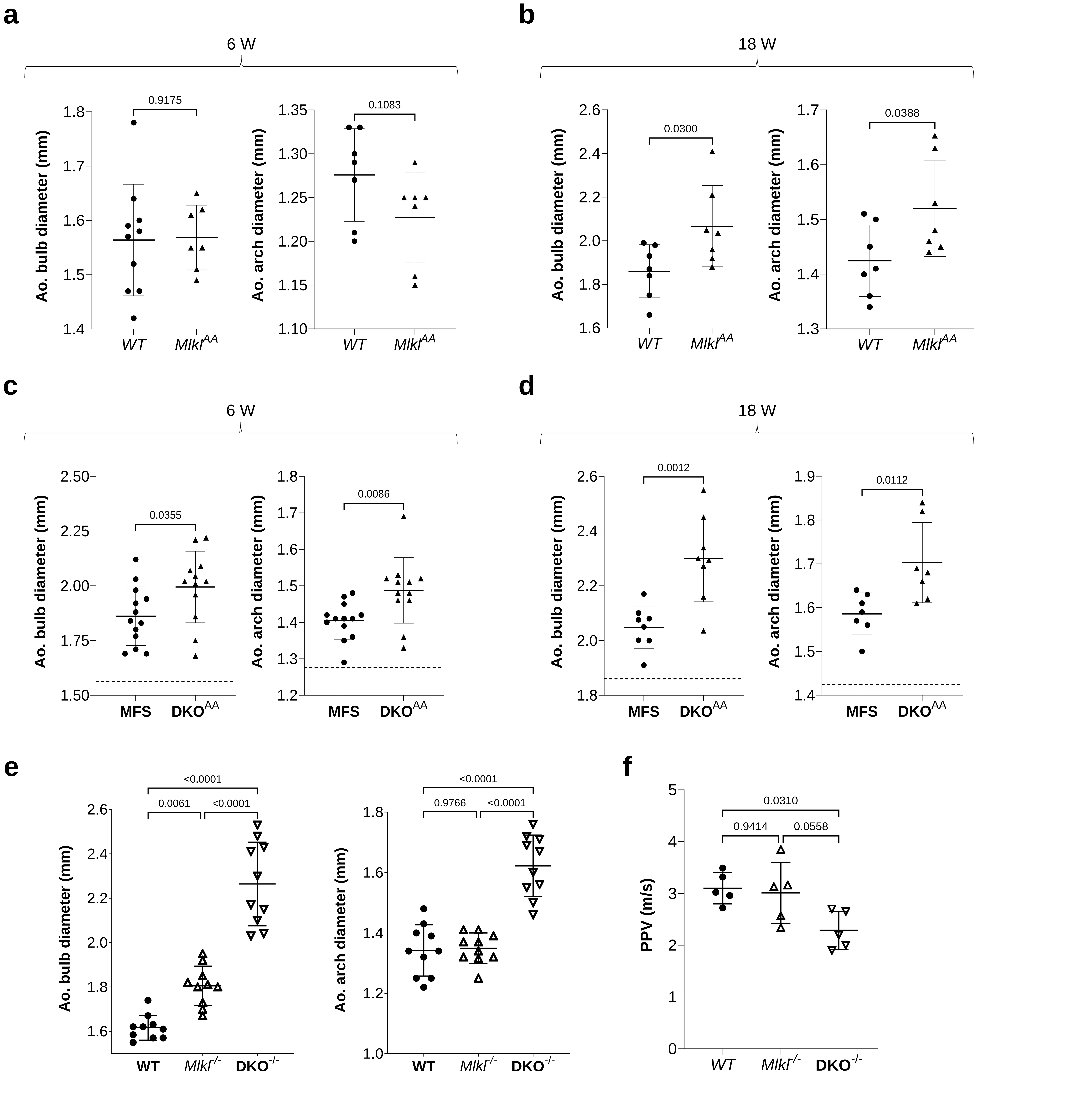


**Supplementary Figure 1. Age-dependent progression of aortic dilatation.**
Related to Figs 1, 2. (a, b) Aortic bulb and arch diameter, WT and *Mlkl*$^{AA}$, at 6-week-old (a) and 18-week-old (b) mice. (c, d) Aortic bulb and arch diameter, MFS and DKO$^{AA}$, at 6-week-old (c) and 18-week-old (d) mice. (e and f) Independent cohort of 12-week-old; WT, *Mlkl*$^{-/-}$ and DKO$^{-/-}$ (*Fbn1*$^{C1039G/+}$; *Mlkl*$^{-/-}$) mice, aortic bulb diameter, and arch diameter (e) and PPV (f). Data are mean ± SD; $n$ = 7 (a, b, c, and d); $n$ = 10 (e); $n$ = 5 (f).

### Supplementary Figure 2

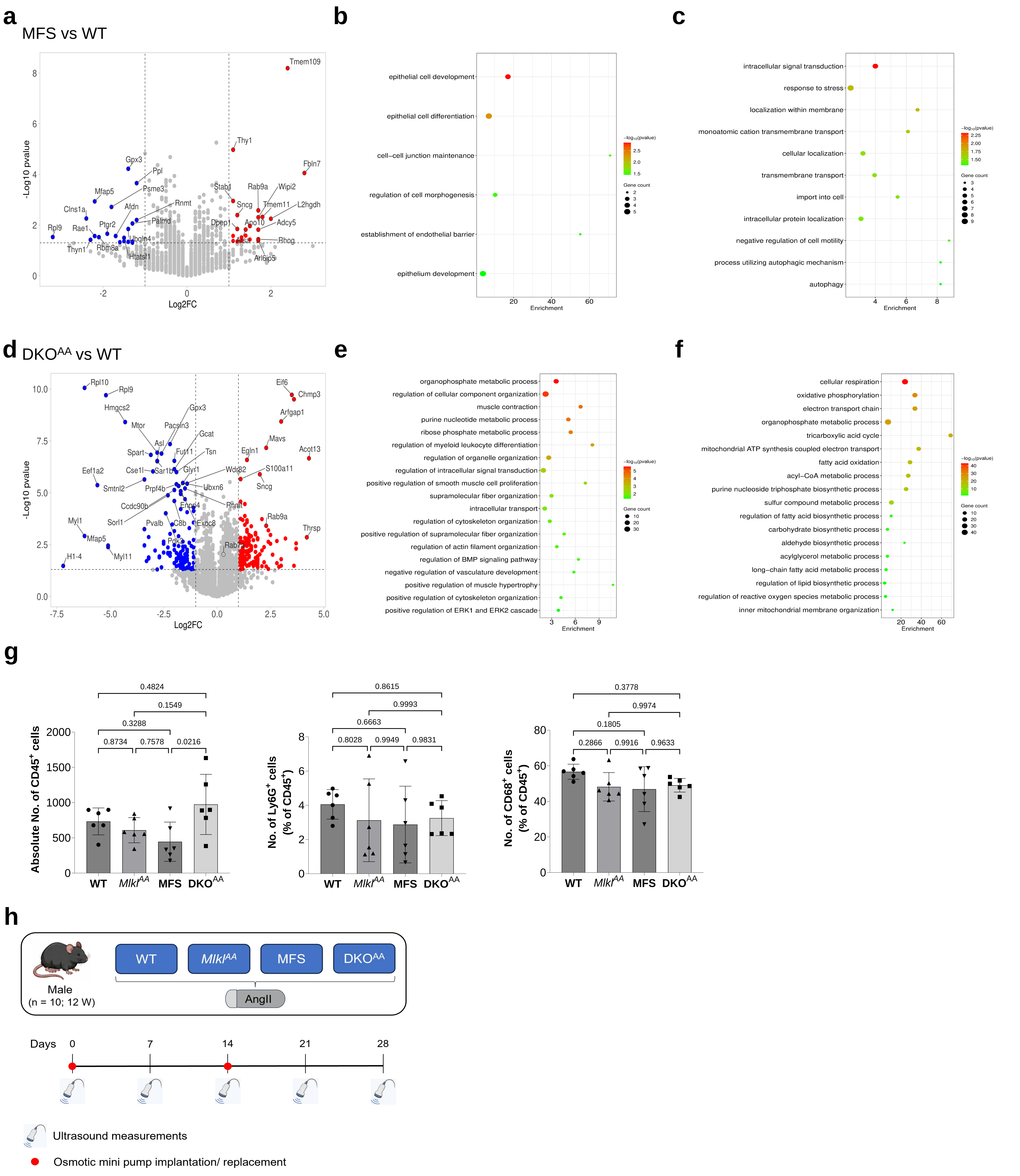


**Supplementary Figure 2. Comparative aortic proteomic remodelling, MFS and DKO$^{AA}$ versus WT, aortic-leukocyte flow cytometry, and AngII study design.**

Related to Figs 2. Global proteomic from MFS versus WT (a) Volcano plot showing most significantly deregulated proteins, (b and c) GO terms enriched analysis from down- (b) and up- (c) regulated proteins in MFS versus WT. Global proteomic from DKO$^{AA}$ versus WT (d) Volcano plot showing most significantly deregulated proteins, (e and f) GO terms enriched analysis from down- (e) and up- (f) regulated proteins in DKO$^{AA}$ versus WT. (g) Flow cytometry analysis of thoracic aortas from 12 week old WT, *Mlkl*$^{AA}$, MFS and DKO$^{AA}$ animals (left) absolute number of CD45$^{+}$ cells (middle) Ly6G$^{+}$ neutrophils and (right) CD68$^{+}$ macrophages as a percent of CD45$^{+}$. (h) Experimental design of the AngII model; WT, *Mlkl*$^{AA}$, MFS and DKO$^{AA}$ males were implanted with AngII osmotic minipumps (replaced on day 14), and followed with ultrasound at days 0, 7, 14, 21 and 28. Data are mean ± SD; *n* = 5 (a-f); *n* = 6 (g).

### Supplementary Figure 3

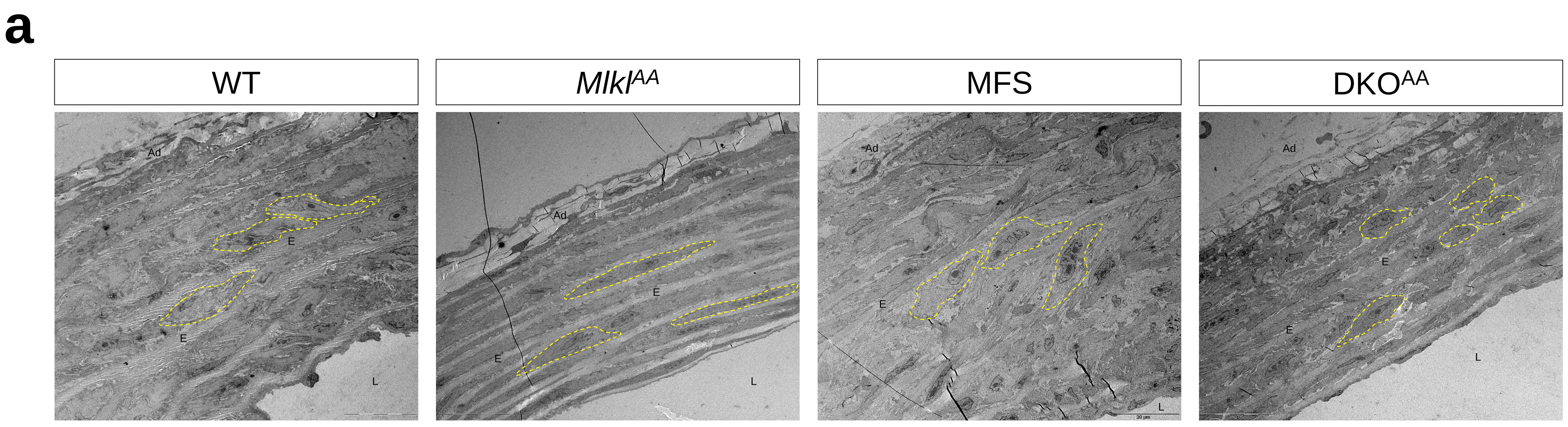


**Supplementary Figure 3. Overview of aortic-wall ultrastructure.**
Related to Figure 4. (a) Representative images of low-magnification TEM survey of the compete aortic wall in WT, *Mlkl*$^{AA}$, MFS and DKO$^{AA}$ animals at 12 weeks. SMC morphology is highlighted with yellow dashed line. Scale bars 20µm. E = Elastin, L = Lumen, Ad = Adventitia. *n* = 1.

Supplementary Figure 4

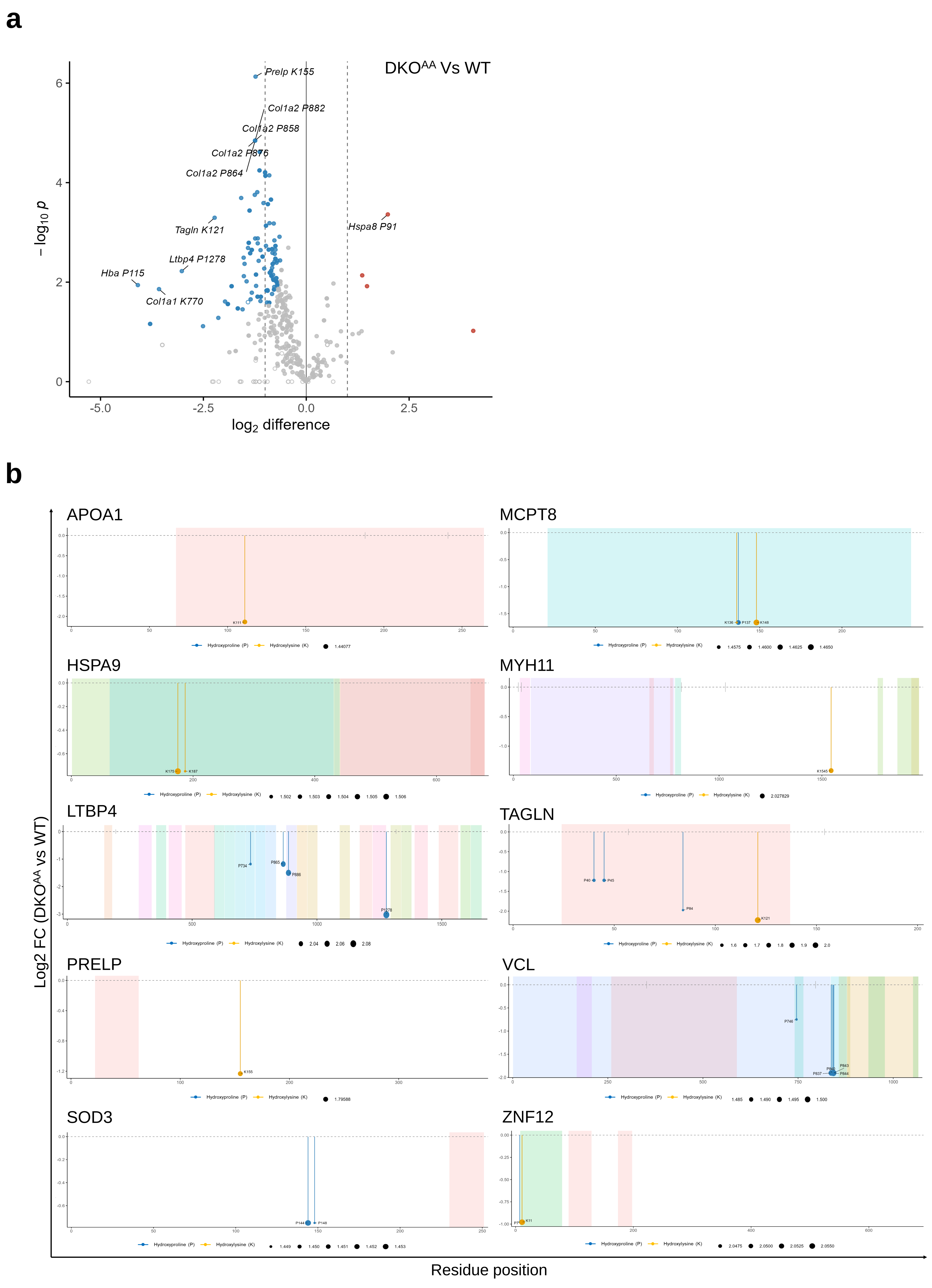


**Supplementary Figure 4. Site-specific hydroxylation changes, DKO$^{AA}$ versus WT.**
Related to Figure 4. (a) Volcano plot of proteins with significantly altered hydroxylation sites. (b) Lollipop plots of significantly altered hydroxylation residues in non-collagen proteins (APOA1, MCPT8, HSPA9, MYH11, LTBP4, TAGLN, PRELP, VCL, SOD3, ZNF12). See Methods for statistical tests.

### Supplementary Figure 5

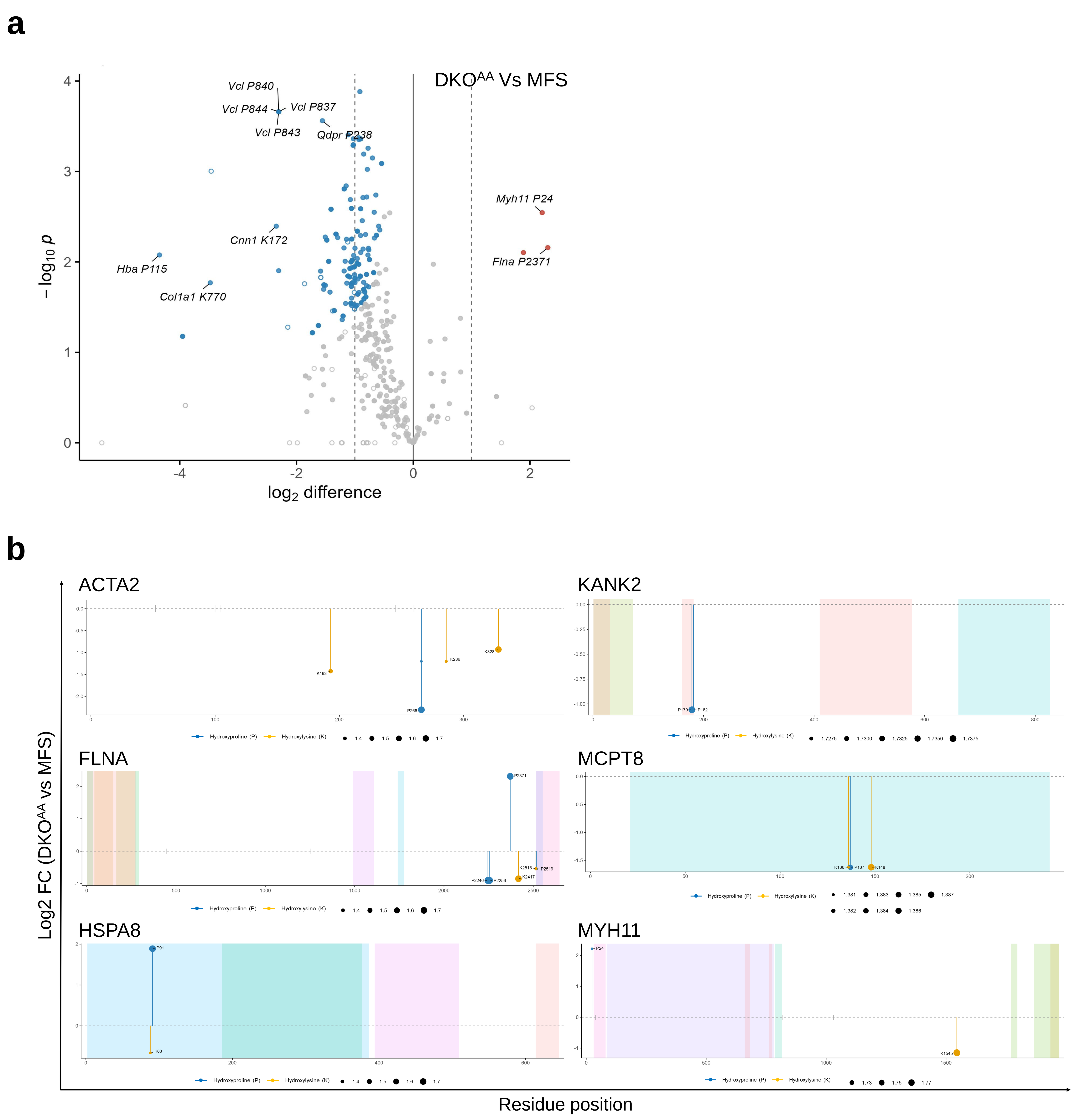


**Supplementary Figure 5. Site-specific hydroxylation changes, DKO$^{AA}$ versus MFS.**
Related to Figure 4. (a) Volcano plot of proteins with significantly altered hydroxylation sites. (b) Lollipop plots of significantly altered hydroxylation residues in non-collagen proteins (ACTA2, KANK2, FLNA, MCPT8, HSPA8, MYH11). See Methods for statistical tests.

### Supplementary Figure 6

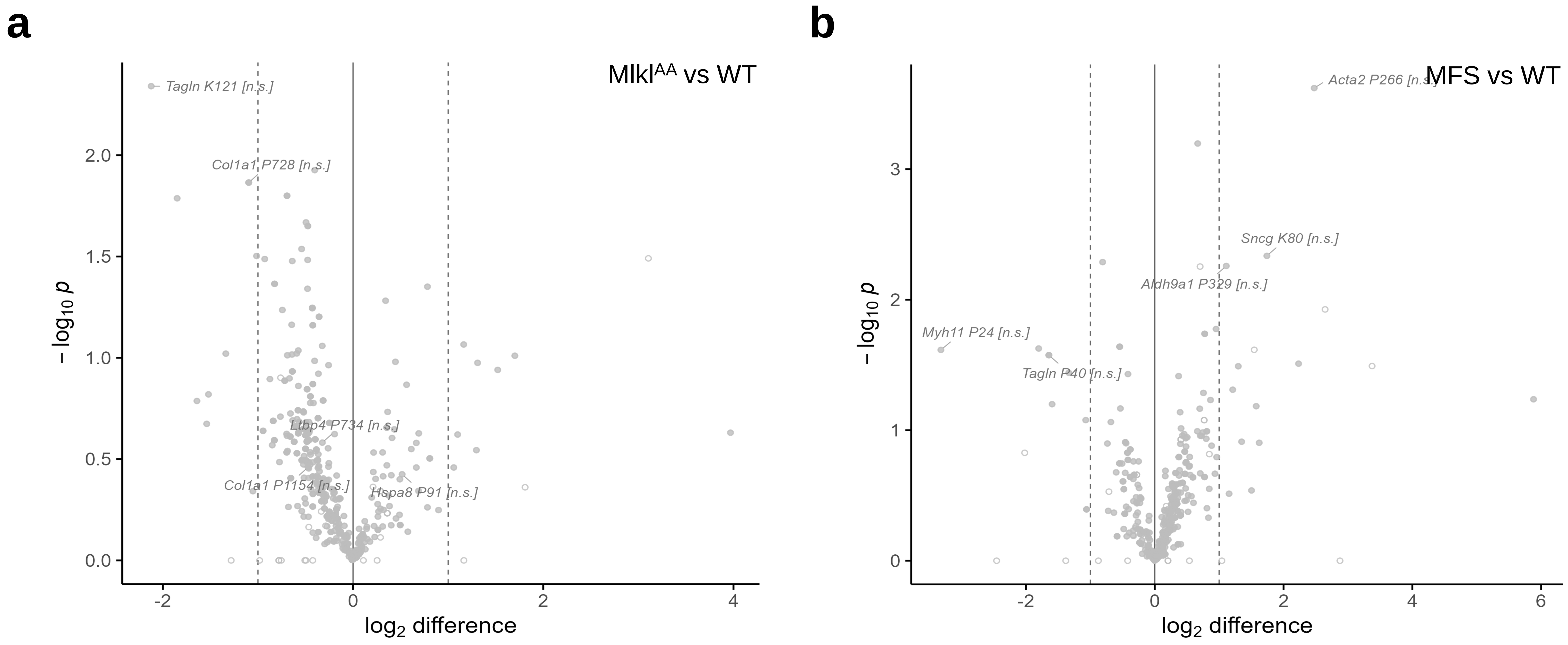


**Supplementary Figure 6. Collagen hydroxylation is unaltered in *Mlkl^AA^* or MFS aortas versus WT.**
Related to Figure 4. (a and b) Volcano plots, *Mlkl^AA^* (a) and MFS (b) versus WT; labelled sites not significant. See Methods for statistical tests.

### Supplementary Figure 7

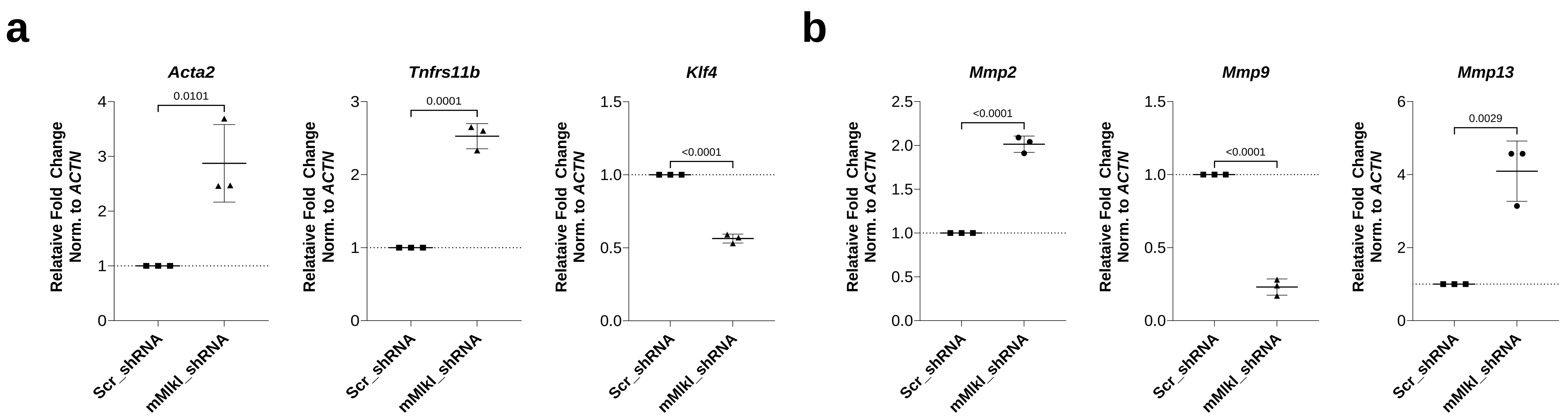


**Supplementary Figure 7. MLKL depletion shifts SMC phenotype markers and MMP expression.**
Related to Figure 5. (a and b) Quantitative RT-PCR in Mlkl-shRNA versus Scr-shRNA mSMCs showing *Acta2*, *Tnfrsf11b* and *Klf4* (a) and *Mmp2*, *Mmp9* and *Mmp13* (b). Data are mean ± SD; *n* = 3 (biological replicates).

Supplementary Figure 8

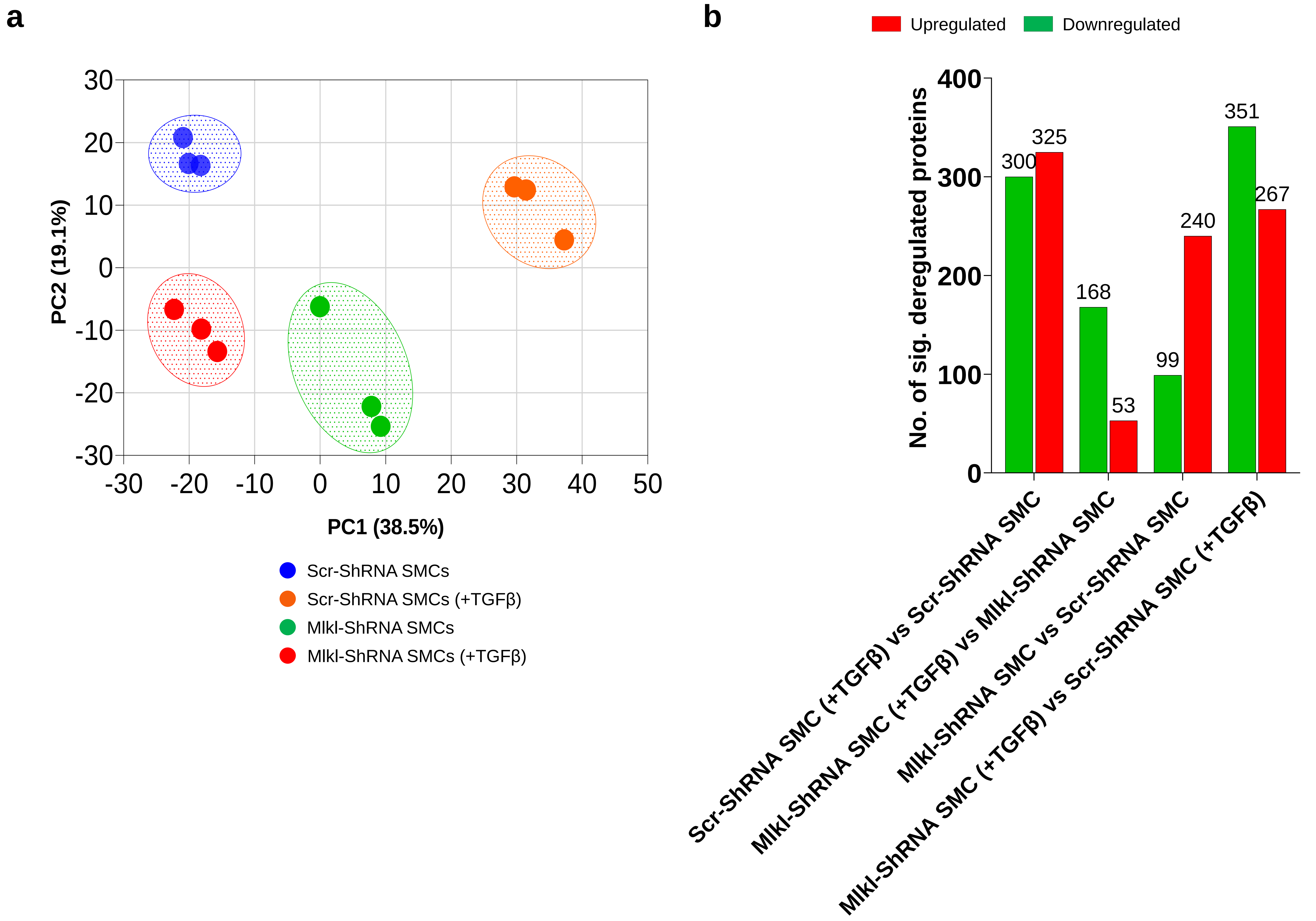


**Supplementary Figure 8. Principal component analysis of the SMC proteome.**
Related to Figure 5. (a) PCA of normalized $\log_2$ LFQ intensities (top 2,000 most variable proteins), (b) Number of significantly deregulated proteins (adj. $p < 0.05$ and $|\log_2 FC| \geq 1$) between each comparison. $n = 3$ (biological replicates colored by group).

### Supplementary Figure 9

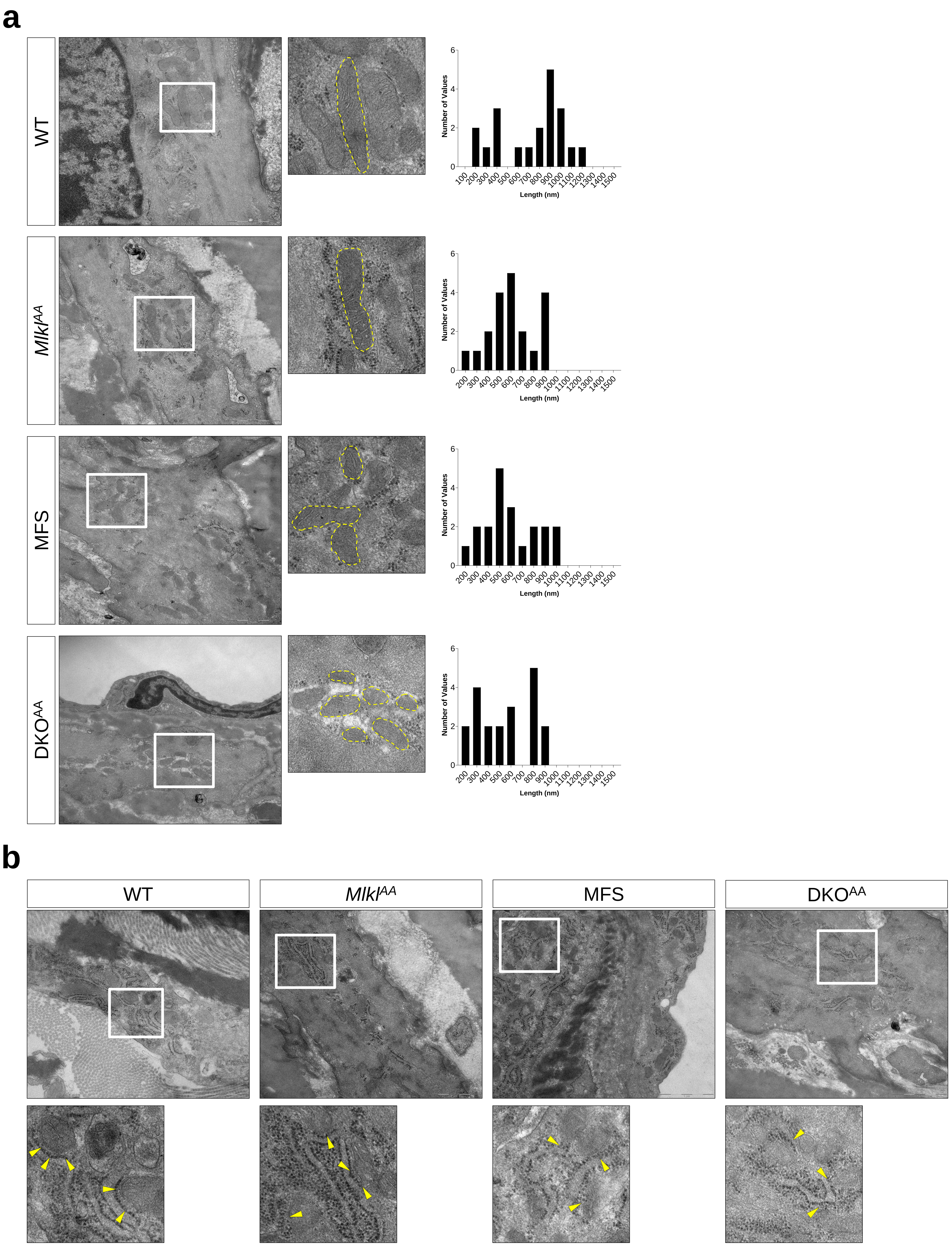


**Supplementary Figure 9. Mitochondrial and endoplasmic-reticulum ultrastructure.**
Related to Figure 6. Representative images of high-magnification TEM survey of WT, *Mlkl*$^{AA}$, MFS and DKO$^{AA}$ animals at 12 weeks. (a) SMC mitochondria (outlined, dashed yellow) with per-genotype length distributions. (b) SMC endoplasmic reticulum with mitochondria–ER contact sites (MERCs) (yellow arrowheads). Scale bars 1µm. *n* = 1.
